# Adaptational lag at high elevations depends on life stage in a California wildflower

**DOI:** 10.1101/2025.09.26.678840

**Authors:** Brandie Quarles-Chidyagwai, Sarah Ashlock, Johanna Schmitt, Julin N. Maloof, Jennifer R. Gremer

## Abstract

1. High elevation populations are expected to receive reduced snowpack, warmer temperatures, and more variable precipitation patterns, potentially putting them at risk if rates of adaptation do not keep pace with climate change. Populations from climates more closely aligned with the changing high elevation conditions may prove better suited to current climate than the current local populations. Thus, it is essential to assess 1) whether high elevation populations are locally adapted to current climate, and 2) whether fitness of lower elevation populations from warmer climates is higher than for local populations in high elevation conditions.
2. We conducted a common garden study with *Streptanthus tortuosus* at a high elevation site. Twenty-three populations from across the species range were measured weekly for mortality, and reproductive output was measured at the end of the growing season. We examined the effects of climatic distance from the site of origin on plant performance. The effects of weekly weather on mortality were also assessed.
3. We observed adaptational lag for high elevation populations, including the native population, but only for some life stages. Low elevation populations had higher survival through the first year and over winter. Additionally, the probability of reproducing was highest for populations from the warmest climates. Warmer ambient temperatures at the high elevation garden were also associated with higher weekly mortality across populations. However, survival to reproduction in the second year was higher in populations from climates closer to the garden, i.e. high elevation populations. Thus, adaptational lag differed among life stages.
4. **Synthesis:** This study highlights the importance of considering variation in life history and seasonal conditions when evaluating how species that occur across an elevational gradient may respond to climate change. This adds to a growing body of evidence that reveals warming temperatures as a threat to high elevation populations. However, unlike previous studies, this threat was not consistent across life stages. These results suggest that strategic assisted gene flow that combines the benefits of warm-adapted low elevation populations with the benefits of snow-adapted life history from high elevation populations may be beneficial in this species, and similar systems.

## Introduction

Climate change is shifting historical climate regimes across the globe (Pachauri et al., 2014; Parmesan & Yohe, 2003). Plants that grow in high elevation, montane environments may be especially at risk of population decline due to warming temperatures, reduced snowpack, earlier snowmelt, precipitation variability, altered biotic interactions, and projected habitat loss (Alexander et al., 2018; Ashfaq et al., 2013; Dettinger et al., 2018; Engler et al., 2011; Fyfe et al., 2017; Halofsky et al., 2021; Mao et al., 2015; Oldfather et al., 2025). Plant species will have to track those changes spatially through dispersal or adapt evolutionarily in order to persist (Aitken et al., 2008). The potential for adaptation depends on many factors, including the standing phenotypic and genetic variation, and the rate of adaptation may not be sufficient to keep pace with the rate of climate change (Aitken et al., 2008; Anderson, 2016). Spatial tracking of ideal climatic conditions will likely require dispersal up in elevation or latitude (Anderson & Wadgymar, 2020; Chen et al., 2011; Fadrique et al., 2018; Kelly & Goulden, 2008; Lenoir et al., 2008; Parmesan & Yohe, 2003; Vitasse et al., 2021). However, even if a species is capable of dispersing to higher elevations, they will likely encounter differences in seasonal timing and temperature and precipitation patterns (Billings, 1974; Gremer, Chiono, et al., 2020). Thus, species may have to shift their life histories in order to persist at higher elevations (Cavieres & Arroyo, 2000; Gremer, Chiono, et al., 2020; Haggerty & Galloway, 2011; Halbritter et al., 2018; Vitasse et al., 2013).

Identifying whether populations are adapted to local climate conditions can facilitate predictions of how they will respond to climate change. Local adaptation is widespread across plant species (Hereford, 2009; Lortie & Hierro, 2022), including across elevational gradients (Anderson et al., 2015; Angert & Schemske, 2005). The implications of such local adaptation are dependent upon the trajectory of climate change. For example, if climatic conditions have already started to shift and the rate and direction of those changes is expected to continue, then finding local adaptation to current climate would suggest that a population may be able to adapt and persist in the face of continuing climate change. However, if climate is expected to shift in a different direction or rate, a population that is adapted to current climate may be at risk due to both maladaptation and reduced genetic variation resulting from recent climate adaptation (Anderson, 2016; Anderson & Song, 2020; Martin et al., 2023). Recent studies have found evidence that populations adapted to historic climate are maladapted to present conditions and may not be able to evolve fast enough to track future changes in climate (Anderson & Wadgymar, 2020; Browne et al., 2019; Villoutreix et al., 2025; Wilczek et al., 2014). Such adaptational lag could increase the risk of population extinction (Aitken et al., 2008; Anderson, 2016; Kopp & Matuszewski, 2014)

Populations adapted to historical environmental conditions similar to the shifting climate in a different part of a species range, may have higher performance than local populations. For example, studies have found that plants from drier (Metz & Tielbörger, 2023) and warmer (Anderson & Wadgymar, 2020; Browne et al., 2019; Martínez-Berdeja et al., 2019; Metz & Tielbörger, 2023; Villoutreix et al., 2025; Wilczek et al., 2014) home sites have performed better than the local population. More broadly, climate distance has been found to be a significant predictor of fitness within and outside species ranges (Montalvo & Ellstrand, 2000; Rutter & Fenster, 2007; Samis et al., 2019; Villoutreix et al., 2025). Dispersal or gene flow from populations adapted to past climates resembling changing conditions could facilitate evolutionary rescue and persistence in populations experiencing adaptational lag (Aitken et al., 2008; Anderson & Wadgymar, 2020; Bontrager & Angert, 2019). In species that occur across elevational gradients, gene flow from low elevation populations may help higher elevation populations to adapt to warming temperatures. Common garden studies with species that grow across elevational gradients can simultaneously test for adaptational lag in high elevation populations and for the potential of low elevation populations to rescue high elevation populations (Anderson et al., 2025; Anderson & Wadgymar, 2020; Kooyers et al., 2019).

Species with populations that occur across an elevational gradient are likely to have life history differences that correspond with the seasonal differences between elevations (Anderson & Wadgymar, 2020; Bucher & Römermann, 2020; Gremer, Chiono, et al., 2020). These life history differences may be more distinct in mediterranean climates where low elevation populations typically have winter growing seasons while high elevation populations grow in the spring and summer after snowmelt (Gremer, Chiono, et al., 2020). This contrast in growing seasons may result in different fitness-climate distance relationships across life stages. For example, low elevation populations may be able to survive during warmer, drier growing seasons better than plants from high elevations, but they may not have the adaptations or plasticity necessary to survive harsh winters, or time germination and reproduction with favorable seasonal conditions. Beyond differences in growing season, there can also be differences in whether plants grow as annuals versus biennials/perennials (Lowry et al., 2008; Menges & Waller, 1983). For example, annual plants may not have sufficient time to grow to reproductive maturity during the short high elevation growing seasons. Furthermore, there may be tradeoffs between first year and second year life stages which may result in maladaptive life history of non-local plants (Cotto et al., 2019; Satyanti et al., 2021). In particular, low elevation plants may grow fast in the first year of life and not have enough remaining resources to survive to reproduce well in the second year of life.

Indeed, seasonal and life history timing are shaped both by growing season conditions, as well as potentially unfavorable conditions that occur throughout the year. Specifically, life history differences may have evolved to both time growth with favorable growing season conditions and to avoid unfavorable conditions in other parts of the year (Giesel, 1976). Thus, fitness components that determine or are directly determined by the seasonal timing of growth, such as germination and over-winter survival, will likely be influenced by climate outside of the growing season more than fitness components not directly related to life history timing. Furthermore, life history differences across elevations can be a form of niche construction, whereby organisms determine the environment that different life stages experience (Donohue, 2003, 2005; Gremer, Chiono, et al., 2020). Such niche construction can reduce the degree of environmental variation experienced, especially during the growing season (Bontrager et al., 2025; Clark et al., 2020; Donohue, 2005). Reduced environmental variation among populations during the growing season due to niche construction may reduce differentiation in growing season traits across elevations, thereby reducing the potential for gene flow from low elevation populations to rescue high elevation populations. That is, if niche construction is no longer conserving the seasonal climate sufficiently to prevent maladaptation at high elevations.

In this study we grew plants from populations across the geographic and elevational range of a native California wildflower, *Streptanthus tortuosus* (mountain jewelflower) in a common garden experiment at high elevation. We examined the effects of climatic distance from the site of origin on plant performance to ask 1) Is the population that is native to the garden site locally adapted to the contemporary garden site climate? Or is it showing signs of maladaptation due to recent climate change?, and 2) Will plants from climates historically similar to the current garden climate, i.e. warmer and drier low-elevation sites, perform better than plants from high elevation sites due to recent climate change? We hypothesized that life history adaptations across the elevational gradient will result in different climate-fitness relationships across the life cycle. We also hypothesized that fitness components at different life cycle stages will respond differently to climate distance between the home site and transplant site based on the full water year versus the growing season.

## Materials and Methods

### Study System

*Streptanthus tortuosus* (Brassicaceae) is an annual, biennial, or short-lived perennial herb that occupies outcrops and dry, rocky slopes throughout California and southern Oregon (Baldwin & Goldman, 2012; Calflora, 2024; Preston, 1991). The species occurs across a broad elevational (200 m to 4100 m) and latitudinal range (from southern California to southern Oregon), resulting in vast differences in seasonal conditions across populations (Figure 1a). Specifically, low-elevation populations receive rain in late fall and winter with warm temperatures, minimal chilling, and no snowpack accumulation (Gremer, Chiono, et al., 2020). In contrast, high-elevation populations receive rain and snow in late fall and winter with cooler temperatures and substantial snowpack (Gremer, Chiono, et al., 2020). Previous studies have identified life history differences among populations that are attuned to the temperature and precipitation differences across the range (Gremer, Chiono, et al., 2020; Gremer, Wilcox, et al., 2020). Specifically, likely due to longer growth seasons at lower elevations, low elevation plants tend to be annual while high elevation plants tend to be biennial or perennial.

**Figure 1.**
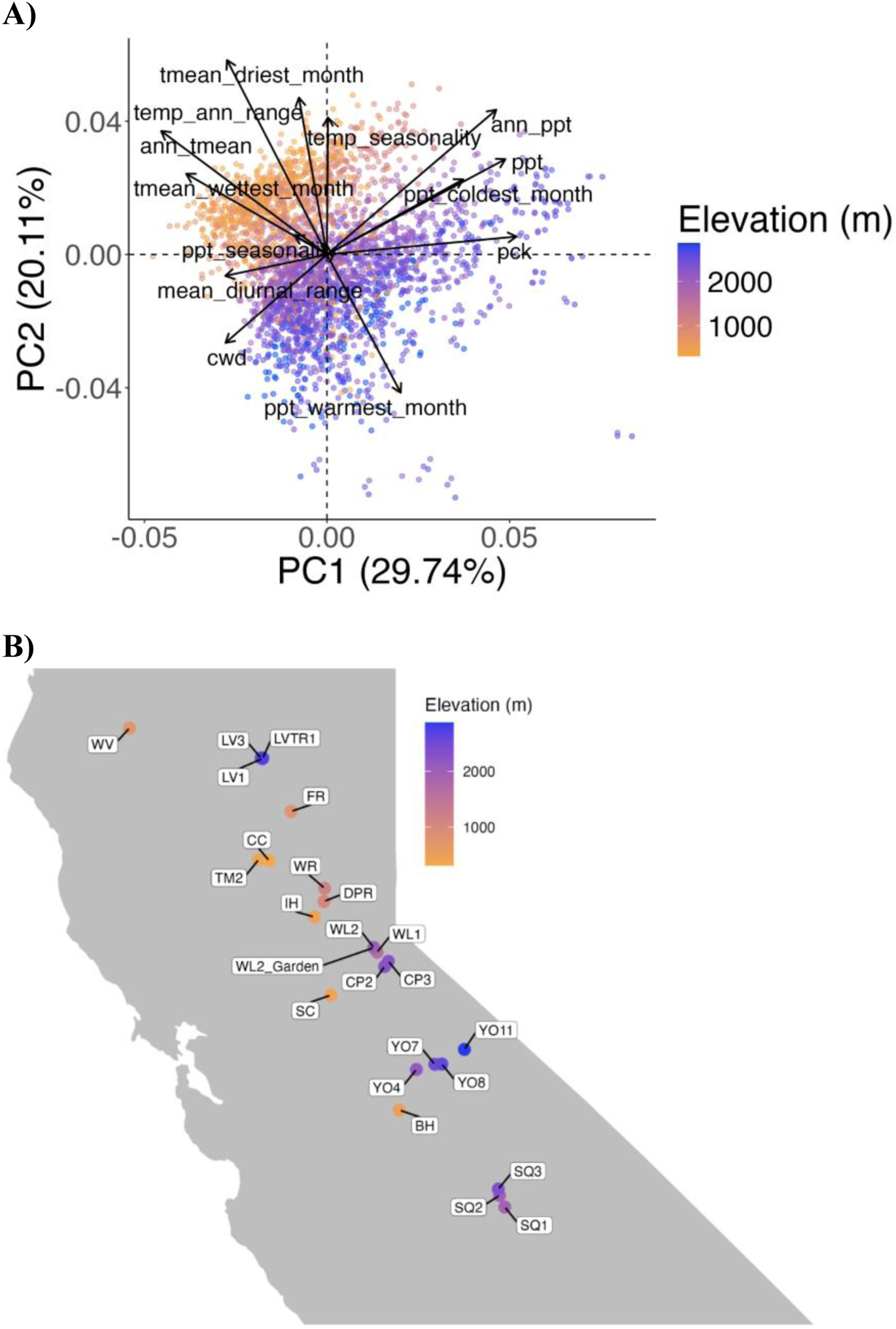
A) PCA showing climate variation across the home sites included in this study. Each point represents a combination of population, time period (recent or historical), and seasonal summary (water year or growth season). PC1 represents a warm/cold (left/right; BIO1), no snow/snow (left/right; PCK) continuum and PC2 represents a temperature annual range (bottom to top: low to high; BIO7), and precipitation in the warmest month (bottom to top: high to low) continuum. B) Map of *S. tortuosus* populations and the common garden site included in the study. Low elevation populations are represented by golden points while higher elevation populations are represented by purple and blue points.

For this study we took seeds from 23 populations across the range (Figure 1b, Table S1). Seeds were collected as maternal seed families from all populations between June and September, depending on the timing of fruit maturation for each population. Seed collection year ranged from 2014 to 2021 due to natural variation in seed availability, as well as occasional limitations in access to sites and personnel (Table S1). We did not collect seed from plants that produced less than five siliques (fruits) or from plants within 1.5 m from a previously sampled plant. Prior to the start of the experiment, seeds were stored dry at room temperature (∼21°C). We used field-collected seeds because our intention was to simulate actual dispersal, in which plant performance could be influenced by maternal environmental effects as well as genotype.

### Common Garden Design

We conducted a common garden experiment at a high elevation site in the Sierra Nevada mountain range in northern California (2020 m a.s.l; 38.82599, -120.2509; Figure 1b). The field site is located near a naturally occurring population of *S. tortuosus* (Wrights Lake 2, WL2) and is in an open area amidst conifer forest on well-drained, granitic soils. Seeds from all 23 populations, with 7 maternal families per population, and 13 replicates per family, were stratified at 4°C in the dark in a cold room at UC Davis. The length of stratification was determined by elevation, with populations from greater than 1600 m getting 8 weeks, and the rest getting 6 weeks. These stratification conditions are based on germination requirements identified in previous experiments (Gremer, Chiono, et al., 2020). On June 6, 2023, after stratification, plants were moved to a growth chamber with inductive conditions with a 12-hour day length to stimulate germination. Plants were initially exposed to 22°C day/8°C night temperatures. Four weeks later, daytime temperatures were increased by 1°C daily (from 24°C to 27°C) to acclimate seedlings to field conditions. Seedlings were transplanted to the common garden site one week later, on July 11-13 and July 19. Due to variable germination rates, the actual sample sizes upon transplant were lower or higher, ranging from 1 to 27 per family, with 2-7 maternal families per population, for a total of 1573 experimental plants (Table S1). Seeds were planted at staggered times to ensure all plants completed stratification and entered germination conditions at the same time. We transplanted seedlings into a randomized block design with 13 blocks spread across 11 beds. Individuals were planted 25 cm apart, based on field observations of average plant distances. All naturally occurring plant species, including *S. tortuosus* plants, within a 5 cm radius of experimental plants were removed to reduce competition and prevent misidentification, but otherwise the naturally occurring vegetation was left intact. It is likely that if the seedlings had germinated *in situ* they might have inhibited the growth of plants within this radius. Plants were watered two to three times in the first week after transplanting to reduce transplant shock, then grown without further irrigation from July 2023 through October 2024.

### Fitness Across the Life Cycle

To determine how fitness varied between populations, survival was monitored every one to two weeks as personnel availability permitted. A census was conducted at the end of the reproductive season on October 27, 2023, in which flowers and fruits were counted for each surviving plant. In 2024, plants were surveyed every one to two weeks for phenology and reproduction and the number of flowers and fruits was recorded for each individual plant when it reached reproductive maturity, i.e. when the plant stopped producing flowers and all the fruits had elongated.

From the field collected survival and fruit data, we computed fitness components at key life stages, including establishment, survival to the end of the first year, survival to budding, winter survival, and fruit production (Figure 2). Establishment was measured as the likelihood of survival for the first three weeks post-transplant. Survival to budding, or the initiation of reproduction, represents survival from establishment to budding in year 1 and survival from the start of the growth season (6/3/2024) to budding in year 2. Survival to the end of the first year represents survival from establishment to the final field survey in October 2023 (10/27/2023). Winter survival represents survival from the final field survey in October 2023 to shortly after snowmelt (6/3/2024). Fruit production was compared only among populations that had individuals survive to budding. For survival to budding and fruit number, years 1 and 2 were analyzed separately. In 2023 and 2024, some plants still had flowers when fruits were counted. Thus, reproductive output based on fruit number might represent an underestimate for later or slower maturing plants. To test how this might have affected our results, we compared an estimate of reproductive output based solely on fruit number to one in which flower and fruit number were summed. We also computed two composite reproductive fitness estimates: probability of successfully reproducing, and total reproductive output. The probability of successfully reproducing is the probability of making at least one fruit in any year. The total reproductive output is the total fruit number of the individuals that successfully made at least one fruit in either year 1 or 2 (or both). Each fitness metric was calculated at the individual level with survival estimates coded as 1or 0 for survival or no survival, respectively.

**Figure 2.**
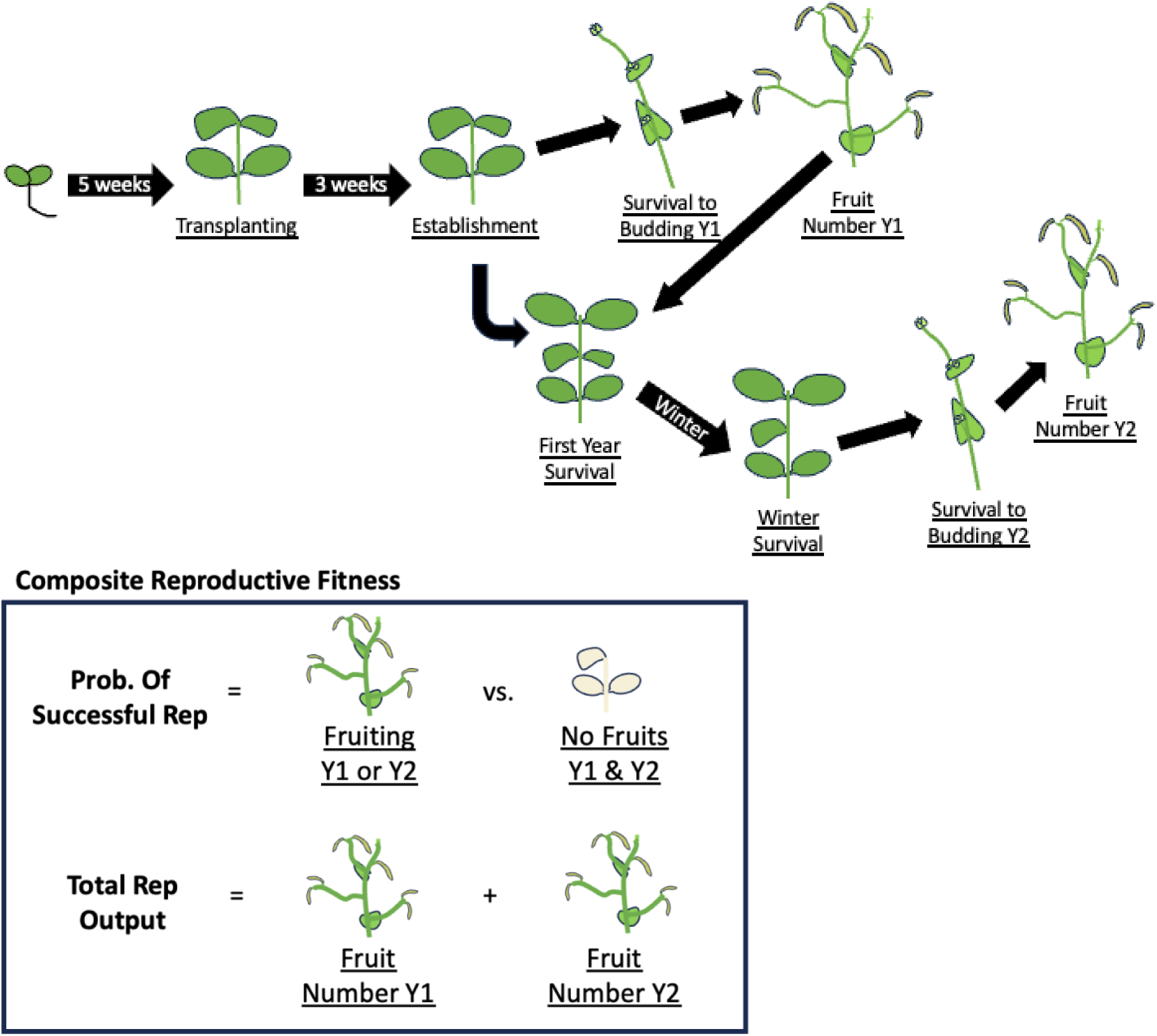
Diagram representing the fitness components measured in this study. Plants were transplanted to the garden when they were 5 weeks old and establishment was measured 3 weeks later. Note that first year survival could be accomplished either by surviving to produce fruits or surviving as a vegetative plant. Probability of successfully reproducing estimated the probability of producing at least one fruit in either year 1 or 2 and total reproductive output was computed as the total number of fruits produced across year 1 and 2.

We tested for tradeoffs between year 1 and year 2 fitness components via linear regressions in R (lm function in the stats package in R; R Core Team, 2024). Regressions were conducted on population means for each vital rate above. Population mean probabilities were arcsine transformed.

### Climate Data

To evaluate whether plant performance depended on the climate of each population’s home site, we obtained recent and historical climate data for each home site and 2023-2024 climate data for the garden site. We classified recent climate as 1994-2023 and historical climate as 1964-1993 for the home sites. We chose for the historical time period to begin in 1964 so that the number of years would be the same for both time periods of climate; other studies have used similar time frames (Browne et al., 2019; Rutter & Fenster, 2007; Samis et al., 2019). All climate data for this analysis were extracted from the Flint Basin Characterization Model (Table 1; Flint et al., 2021). In addition, we used those ‘Flint’ variables to calculate six ‘bioclim’ climatic variables (Table 1; *Bioclimatic Variables — WorldClim 1 Documentation*, n.d.) and four ‘bioclim-like’ variables that capture temperature and precipitation in extreme times.

**Table 1.**
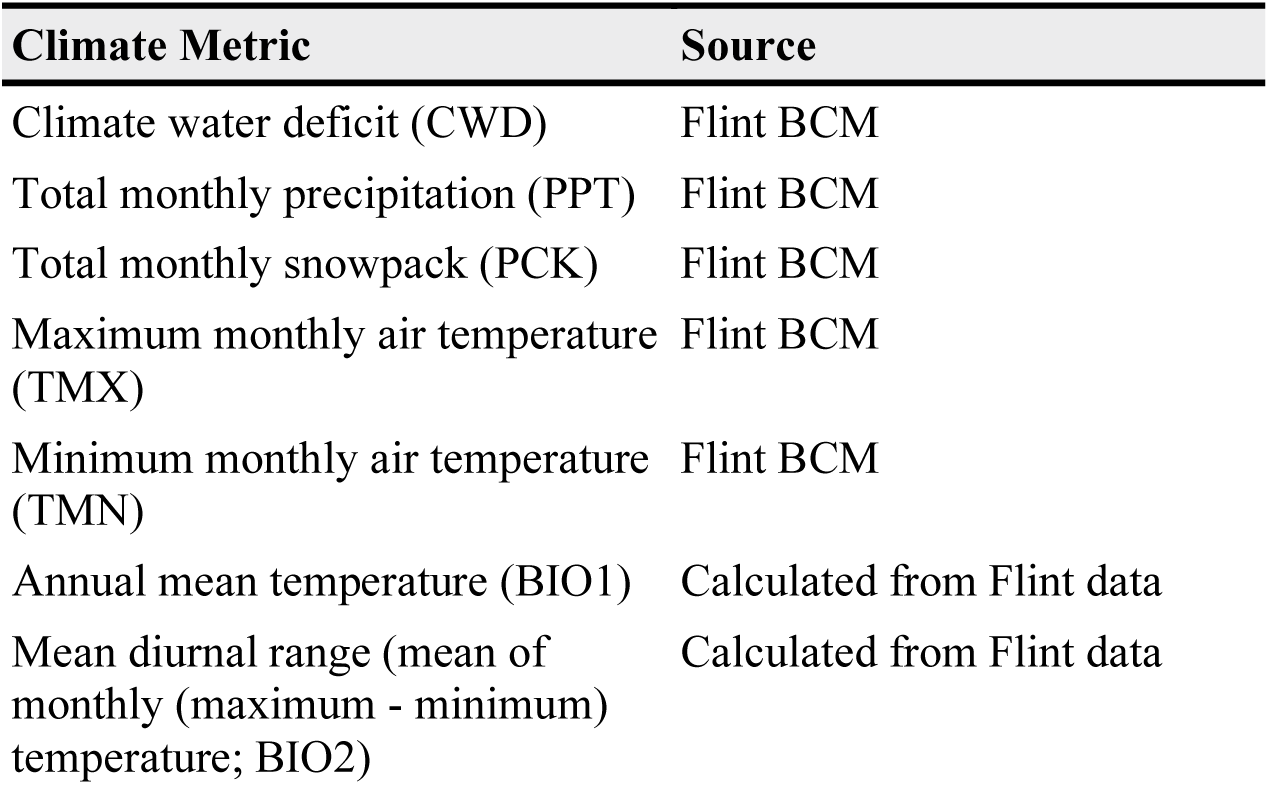
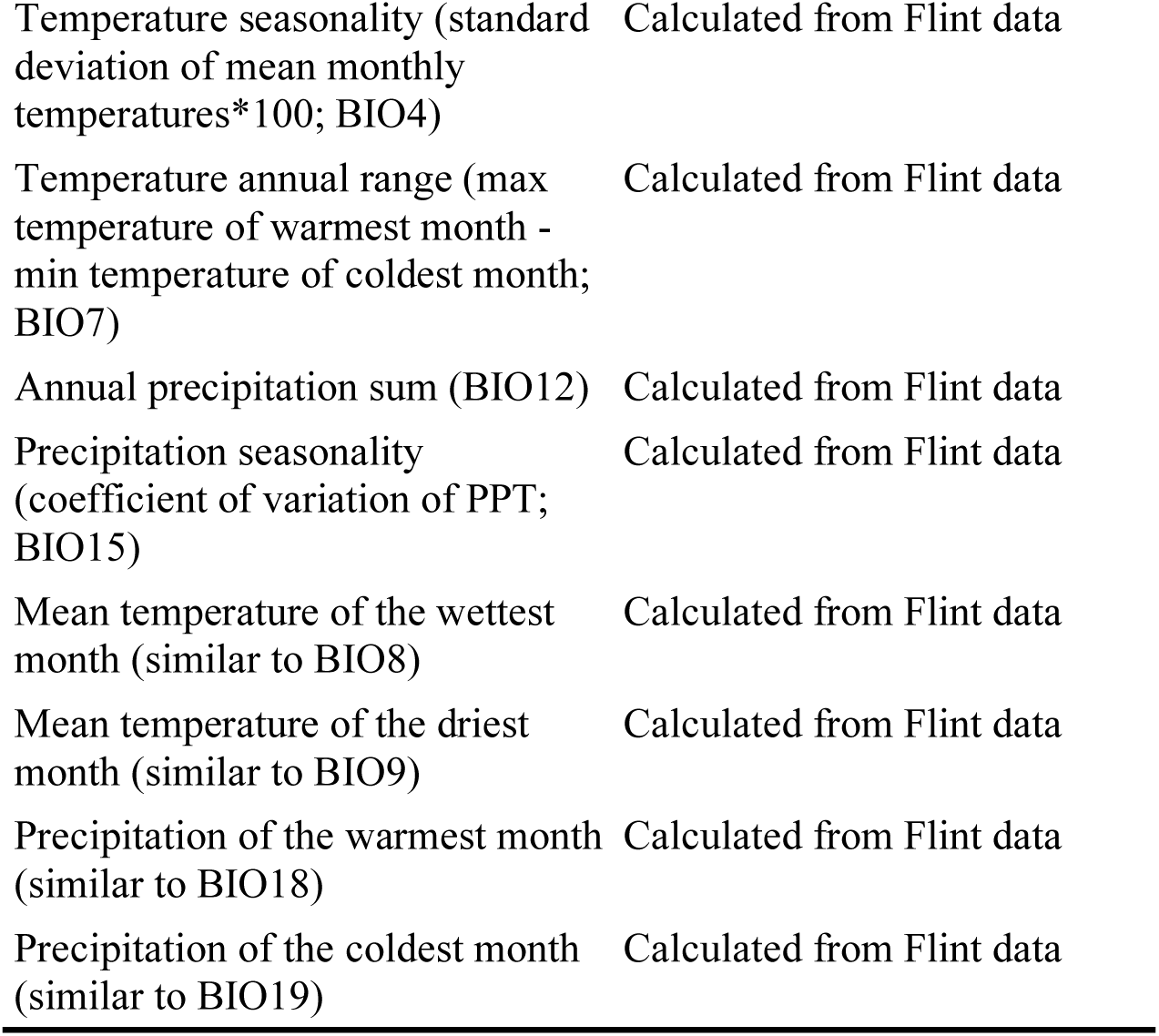
Climate metrics used in this study and the source of the data. We used five ‘Flint’ variables, six ‘Bioclim’ variables calculated from the Flint variables, and for ‘bioclim-like’ variables calculated from the Flint variables. CWD represents the annual evaporative demand that exceeds available water and is an estimate of drought.

#### Seasonal Summaries of Climate

We summarized the climate data in two ways, by water year (November through October) and by the estimated growth season for each population. We set the water year to end with October, rather than start in October as other studies do, to match the timing of our end of year fitness measurements. We estimated the growth season for each population based on how much snow they typically received from 1964-2023. For populations that have received, on average, less than 70mm of snowpack a year (Table S1), the first month of the growth season was calculated as the average month where PPT was greater than or equal to 25 mm, an amount likely to stimulate germination in this species and other annual plant systems (Beatley, 1974; Gremer, Chiono, et al., 2020; Schwinning & Sala, 2004; Tevis, 1958; Went, 1949), and the last month was the month where CWD was greater than the 60-year 3rd quartile of CWD for that group of populations. For populations that, on average, received greater than 70 mm of snowpack a year, the first month of the growth season is the month where PCK was 0 and TMN was greater than 0 while the last month was the month where PCK was greater than 70mm, or PCK was greater than 0 and TMN was less than 0, or when TMN was less than or equal to -5C which is equivalent to a moderate freeze (Pardee et al., 2018). In other words, the beginning of the growing season started when the snowpack had melted, and warm temperatures facilitated growth and ended when the snowpack and freezing temperatures returned. We chose a threshold of 70mm for the end of the season based on the amount that would cover the mean height of living plants at the WL2 garden (70 mm in October 2023). This quantitative method of defining the growth season resulted in growing months that match previous observations of the natural populations (Gremer et al., 2020; Gremer *unpublished*). Note that the first month of the growth season as described above was excluded from the final growth season months because that month represents germination and we germinated plants in growth chambers, not the field. Thus, we were able to compare the growth season that the plants experienced in the field to the average growth seasons they received at their home site.

Growth season months were determined for each year separately, allowing for variation in growing season length over time. Some populations and years did not have a month that met the end of the growth season criteria outlined above. For those cases, we used the maximum “last month” across all the other years for each population. Finally, for the BH population in 1979, there was high CWD in the second month of the growing season, which likely represented a mid-season drought. Rather than have a one-month growth period, we used the next month with high CWD as the last month.

### Distance from Home Site

To compare fitness across populations from home sites more or less similar to the garden, climate distance was calculated for each time period (recent and historical) and season (water year and growth season). While we paused measuring the plants in October 2023 and 2024, the final month of the growth season was estimated the same way as the home climates. We evaluated the effect of directional climate distance in annual mean temperature (BIO1) and annual precipitation sum (BIO12) by subtracting the garden temperature or precipitation from the home temperature or precipitation. Temperature and precipitation both vary across elevation (Billings, 1974), and may shift with climate change (Portmann et al., 2009), potentially at different rates across elevation (Pepin et al., 2022). BIO1 and BIO12 were also highly correlated with the other temperature and precipitation variables, respectively (Figure S1).

Since climate is ultimately multivariate, we also calculated the multidimensional absolute climate distance between the garden site and the recent and historical climates of each home site. We used Gower’s similarity metric (Gower, 1971) as described in (Rutter & Fenster, 2007):

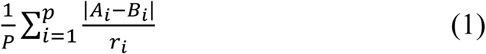

The number of climate variables is represented by *p*, A_i_ and B_i_ are the climatic values with A as the garden site and B as the home site, and r_i_ represents the total range of the climatic variable in the dataset. Bootstrap 95% confidence intervals were calculated for each population’s climate distance estimates using the boot package (Canty et al., 2024). The bootstrap sampled each year of climate data, with replacement. For each bootstrap replicate, the recent and historical 30-year averages of each climate variable were determined and then used to calculate the Gower’s distance. The function boot.ci was used with the “norm” method (Carpenter & Bithell, 2000).

We calculated the geographic distance between each home site and the garden site from their longitudes and latitudes (Table S1) with the haversine formula implemented in R version (4.4.1) using the distHaversine function in the geosphere package (Hijmans et al., 2024). Elevation distance was calculated by subtracting the garden’s elevation from each home site’s elevation.

We tested whether Gower’s climate distance of each population from the garden site was correlated with geographic distance and elevation distance with a Pearson’s correlation test in R (cor and cor.mtest functions in the corrplot package; Wei et al., 2024).

### Relating Fitness to Climate and Geographic Distance

To test whether plant fitness at the common garden depended on the climate distance and/or geographic distance between the home sites and the garden site, we fit mixed effect models with each individual fitness component, and the composite reproductive fitness estimates as dependent variables. Climate distance (temperature, precipitation, or Gower’s) was a fixed effect while population and block were included as random effects. Geographic distance was not significantly correlated with climate distance or elevation distance, allowing it to be included as an additional fixed effect in all models. Maternal family nested within population was also included as a random effect when sample size permitted and it explained greater than zero variation in fitness. Separate models were tested for each season and time period. The recent and historical water years of the home climates were compared to the 2023 water year (Nov 2022-October 2023) for year 1 fitness metrics (establishment, first year survival, survival to budding and fruit number in year 1) and the 2024 water year (Nov 2023-October 2024) for year 2 fitness metrics (over-winter survival, survival to budding and fruit number in year 2). The recent and historical growth seasons of the home climates were compared to July-December 2023 and June-November 2024 at the garden. Populations with data from fewer than 3 individuals were removed from the models for each fitness trait.

We analyzed the number of fruits in year 2 and total reproductive output (log transformed to improve normality) using generalized linear mixed models with population and block as random effects. The average of the 2023 and 2024 climate distance was used in the total reproductive output models which included fruits produced in both years. Maternal family nested within population was included as a random effect for fruit number in year 2. We did not analyze year 1 fruit number since only TM2 individuals produced fruits that year; year 1 fruit number was still included in the analysis of total reproductive output.

All survival components of fitness (establishment, survival to budding, first year survival, and winter survival) and the composite probability of successfully reproducing were analyzed using mixed effects logistic regressions (glmer function in the lme4 package in R; Bates et al., 2015). Maternal family nested within population was included for first year survival, over-winter survival, and the probability of successfully reproducing. As with total reproductive output, the average of the 2023 and 2024 climate distance was used for modelling the probability of successfully reproducing. The model with historical growth season temperature and precipitation climate distance did not converge for the probability of successfully reproducing; the results are not reported for that model. Only the effect of the water year climate distance was tested for winter survival since the growth season does not include the winter. Survival to budding in year 1 was not analyzed due to only two populations (TM2 and FR) transitioning to that stage in year 1.

### Weekly Weather during the Common Garden

To complement the climate distance results and evaluate the effects of within season weather events, we measured soil temperature and moisture during the first year of the garden. Specifically, soil temperature (DS1921G-F5 Thermochron iButton; Maxim Integrated Products, Inc) and soil volumetric water content (EC5 Soil Moisture Sensors; METER Group, Inc.) were recorded hourly. There were 5 total temperature sensors, with two in one of the beds and three in another bed; sensors were distributed across each bed. There were 5 soil moisture probes distributed across one bed. We calculated a daily average, maximum, and minimum soil temperature and moisture. Diurnal range was calculated as the daily maximum soil temperature minus the daily minimum soil temperature. Negative VWC values resulted from air pockets developing in very dry soil, so all negative VWC values were converted to zero before calculating the daily average, max, and min (Campbell, 2019).

For all analyses we computed the average, minimum, and maximum, soil temperature and moisture for the week or two weeks before each survival census, from the daily average data. We made the same calculations for the first week of the two-week interval (days 6-13 prior to census). The average diurnal range for the same time intervals was also calculated. It is important to note that soil temperature spanned from the third day of transplanting (7/13/2023) to 10/27/2023 and soil moisture spanned from the day after transplanting was completed (7/20/2023) to 10/20/2023.

### Relating Mortality to Weekly Weather

To test for the effects of weekly soil temperature and moisture on weekly mortality, we tested a mixed effect logistic regression with weekly mortality as the dependent variable, the number of weeks since transplant and soil temperature or moisture as fixed effects, and parent population and block as random effects. A separate model was conducted for each weather summary (min, max, mean, and diurnal range) and time interval prior to each census.

### Estimate of Climate Change at Home Sites

In order to estimate how much climate change the populations have already experienced at their home sites, we evaluated the distance between the historical and recent climate. A PCA was computed for the average climate of all home sites with water year and growth season and each time period included (Table S2). TMN and TMX were removed from the PCA as they were highly correlated with annual mean temperature (Figure S1). To test for significant differences between historical and recent climate, we performed a permanova with season, time period, elevation, and latitude on the euclidean distance between all points in PC space. To follow up on the results from the permanova we then fit a generalized linear model with season, time period, elevation, and latitude for each PC axis (glm function in the stats package in R). We used the predict function in R to determine where the 2023 and 2024 garden climate fell with respect to the home sites’ climate space. For further contextualization of the 2023 and 2024 garden climate we also constructed PCAs with the yearly climate averages for all populations and the garden site. As in the ‘climate change’ PCA, TMN and TMX were excluded.

## Results

### Climate change at the home sites

If climate change has already been impacting *S. tortuosus* populations, then the recent high elevation climate would be more similar to a historical low elevation climate. All populations in this study have experienced a change in climate from the historical climate (1964-1993) to the recent climate (1994-2023) in both the water year and growth season (F=10.56, P=0.001; Figure 3; Table S3). Both the water year and growth season climate shifted towards a warmer climate with less snow and higher annual ranges in temperature. High elevation populations have shifted to be more similar to historical low elevation climates in terms of temperature. Climate at low elevation has shifted less than climate at high elevation (Time period x Elevation for PC1: F=5.78, P=0.019).

**Figure 3.**
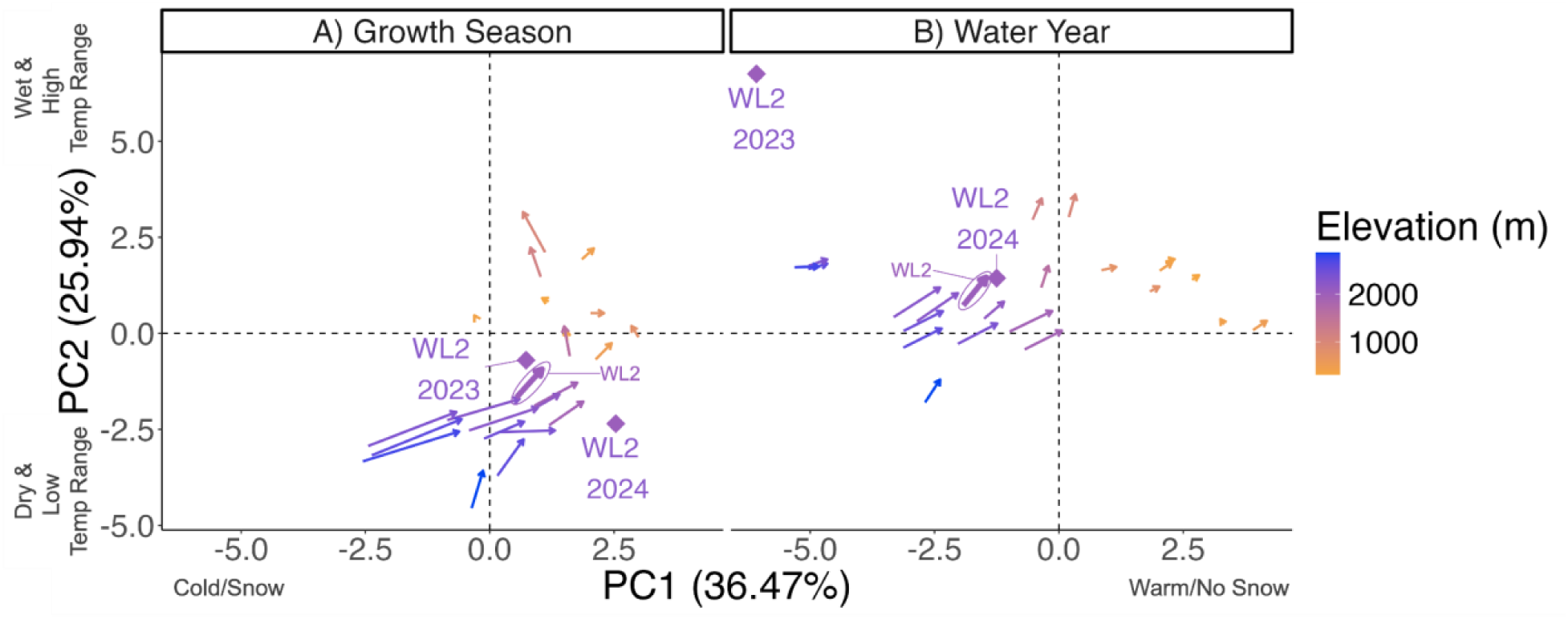
PCA showing climate change at the home sites for the growth season (A) and the water year (B). Arrows show the change from historical (1964-1993) to recent (1994-2023) climate means at each population home site. The line for the shift in climate at the WL2 population is circled and labeled. Diamonds represent the garden climate in 2023 and 2024. PC1 represents a cold/warm (left/right; BIO1), snow/no snow (left/right; PCK) continuum while PC2 represents a dry/wet (bottom/top; CWD) and temperature annual range (bottom to top: low to high; BIO7) continuum (see Table S2). There are significant shifts in PC space from historical to recent climate (Table S3).

### Characteristics of the home sites and garden’s climate

The garden climate mostly fell within the range of the 30-year recent and historic climate averages for the home sites (Figure 3). When it fell outside the home sites’ climate space, the deviation aligned with the direction of climate change observed at those sites. The 2023 garden climate represented a wetter, snowier water year than other years at that site (Figure S2). In contrast, the 2024 and 2023 garden was in the same climate space as the recent 30-year averages at WL2 for the water year and growth season, respectively. The 2024 garden growth season was warmer than other growth seasons at that site (Figure 3a), but fell within the typical yearly variation in temperature across the high elevation sites (Figure S2). Of note, the recent growth season was more constrained than the recent water year climate for the home sites (Figure S2). This trend is not repeated with the historical time period, potentially due to summer precipitation events expanding the climate space of the growth season.

The populations in this study varied in their climate distance from the common garden during the water year and growth season, recent and historic time periods (Figure S3). In addition, both recent and historic water year climate distances were negatively correlated with elevation distance (Recent: r=-0.84, P<0.0001; Historic: r=-0.63, P=0.001). Thus, the greater the absolute climate distance, the more negative the elevation distance; in other words, low elevation populations had higher absolute climate distances from the garden. There were no significant correlations between growth season climate distance and elevation distance. Neither the historic nor recent climate at low elevations resembled the 2023-2024 conditions at the garden, with respect to absolute climate distance.

### Do populations from climates more similar to the garden perform better?

Lower elevation populations from warmer and drier climates than the garden performed better than populations from climates more similar or colder than the garden (Figure 4; Table 2). Establishment, first year survival, and probability of successfully reproducing all increased with increasing temperature distance in the water year (Figure 4a, S4a-b). Establishment also increased with increasing temperature distance in the growth season (Figure S5a) Therefore, populations from warmer climates than the garden had higher first year fitness. In addition, populations from drier environments than the garden had a higher probability of establishment (Figure S4c). However, survival to budding in year 2 and total reproductive output were lower in populations from recently warmer water year and wetter growth season climates, respectively, than the common garden (Figure 4b, S5b). Populations from warmer growing seasons had higher total reproductive output, but this relationship is only positive and significant after taking into account the negative relationship with precipitation distance (Figure S5b). Note that only TM2, a low elevation population, successfully reproduced in both years of the garden (Figure 4).

**Figure 4.**
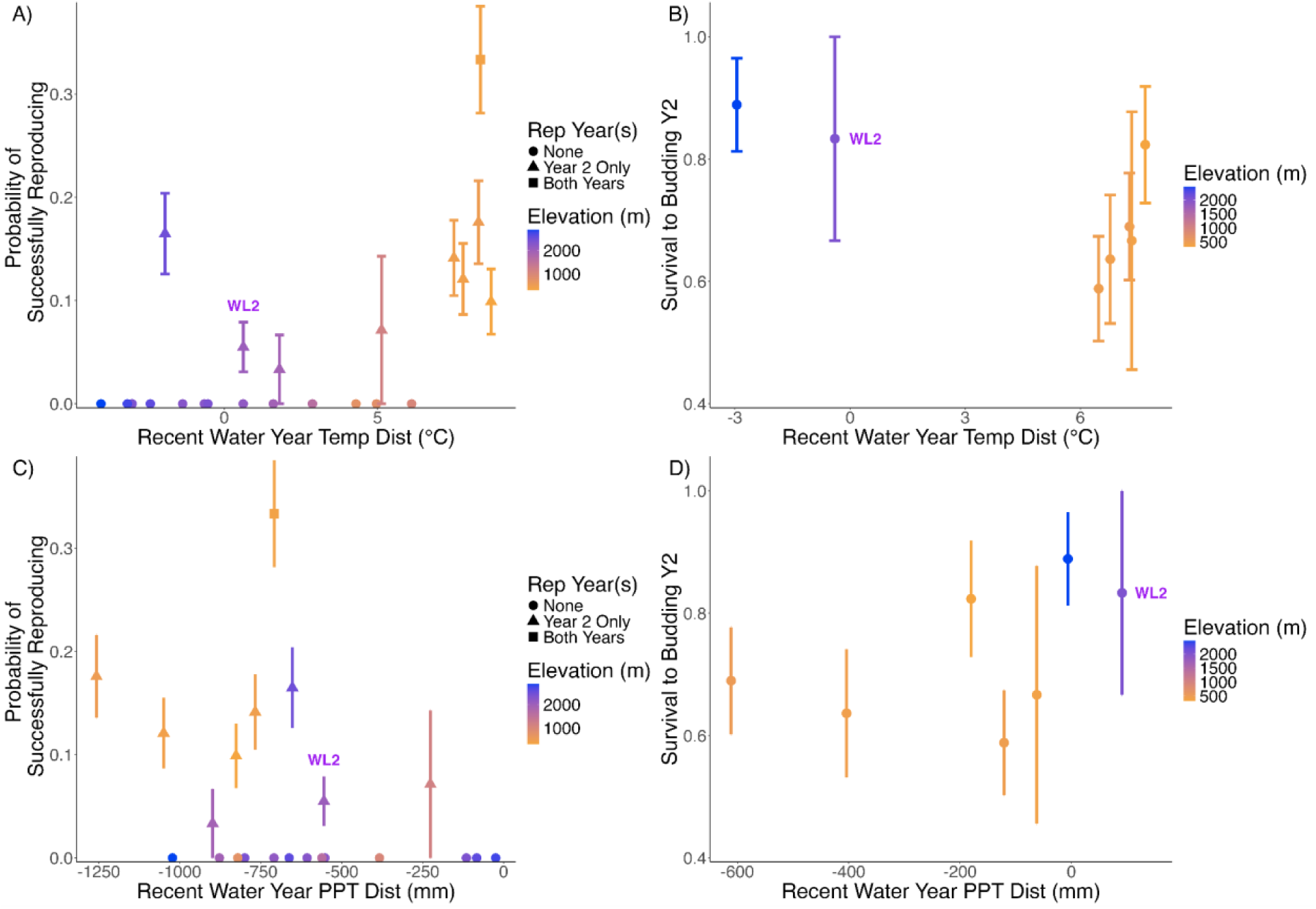
The relationship between (A-B) recent water year directional annual mean temperature (BIO1) distance from home, or (C-D) recent water year annual precipitation sum (BIO12) distance from home and (A,C) the probability of successfully reproducing in either year 1 or 2, and (B,D) survival to budding in year 2. In (A,C) the years in which a population had reproduction are represented by shapes. In all panels, positive values for temperature distance represent warmer temperatures at home than at the garden site and negative values for precipitation distance represent drier home sites than the garden. Points represent population means +/- standard error. Colors represent population elevation in meters. The native WL2 population is labeled.

**Table 2.** Results from linear and logistic regression testing for an effect of the directional climate distance of annual mean temperature (BIO1) and annual precipitation sum (BIO12) from the garden on fitness. Individual fitness components are shown, as well as two composite reproductive traits: probability of successfully reproducing (Prob. of Fruits) and total reproductive output. Establishment, survival to budding in year 2, first year survival, winter survival, and probability of reproduction were analyzed using logistic regressions; all other traits were analyzed using linear regressions. Geographic distance was included in the models as a fixed effect with population and block as random effects. Maternal family nested within population was also included as a random effect in the first year and winter survival, and fruit number in year 2 models. Only 2 populations had greater than 1 individual survive to reproduce in year 1 so we did not analyze survival to budding and fruit number for year 1. For each trait, the effects of water year vs. growth season and recent vs. historical climate were analyzed separately, resulting in 4 models. Only the effect of the water year climate distance was tested for winter survival since the growth season does not include the winter. The two composite reproductive fitness estimates were analyzed with the average climate distance from the garden across 2023 and 2024. The model for probability of reproduction did not converge for historic growth season; the results are not shown. Coefficient estimates are shown with asterisks representing significance. P-values are represented as ***P < 0.001, **P < 0.01, *P < 0.05, ^P <0.08.

|  | Model 1 - Water Year, Recent |  |  | Model 2 - Water Year, Historical |  |  | Model 3 - Growth Season, Recent |  |  | Model 4 - Growth Season, Historical |  |  |
| --- | --- | --- | --- | --- | --- | --- | --- | --- | --- | --- | --- | --- |
|  | Temp Dist | PPT Dist | Geo Dist | Temp Dist | PPT Dist | Geo Dist | Temp Dist | PPT Dist | Geo Dist | Temp Dist | PPT Dist | Geo Dist |
| Estab. Surv | 0.29*** | -0.15* | -0.25*** | 0.28*** | -0.16* | -0.24*** | 0.42** | -0.14 | -0.24** | 0.36** | -0.06 | -0.21* |
| First Year Surv | 1.00*** | -0.36 | -0.08 | 0.99*** | -0.35 | -0.04 | 0.85^ | -0.02 | -0.04 | 0.73^ | 0.18 | 0.02 |
| Winter Surv | 1.04 | -0.42 | -0.03 | 1.00 | -0.45 | 0.02 | N/A | N/A | N/A | N/A | N/A | N/A |
| Surv to Buds Y2 | -0.66* | 0.04 | 0.32 | -0.67^ | 0.02 | 0.28 | -0.86^ | 0.26 | 0.15 | -0.85^ | 0.23 | 0.10 |
| Fruit N Year 2 | -0.14 | -0.09 | 0.19 | -0.15 | -0.10 | 0.19 | -0.05 | -0.08 | 0.19 | -0.04 | -0.09 | 0.19 |
| Prob. of Fruits | 1.80* | -0.59 | -0.31 | 1.76* | -0.59 | -0.24 | 1.31 | 0.42 | -0.17 | N/A | N/A | N/A |
| Total Rep Output | -0.38 | -0.42 | -0.01 | -0.42 | -0.43 | 0.00 | 0.31* | -0.63*** | 0.04 | 0.32* | -0.63*** | 0.08 |

To take into account the multivariate nature of climate, we also tested for the effects of multidimensional climate distance. Similar to temperature climate distance, Gower’s climate distance from home climate was a significant predictor of fitness, and the direction of the relationship differed across fitness components (Table 3). Establishment, first year survival, winter survival, and probability of successfully reproducing all increased with increasing recent water year climate distances from the garden (Figure 5a, S6). Similarly, there was a positive relationship with the historic water year climate distance for first year and winter survival. In other words, populations with climates more different from the garden climate had higher fitness in year 1 and over the first winter. In contrast, populations from historic home climates more similar to the garden climate had higher establishment and first year survival (Figure S7a-b). In concordance with the temperature distance effects, survival to budding in year 2 decreased with increasing recent and historic water year and recent growth season climate distance (Figure 5b, S7c). Total reproductive output also decreased with increasing recent and historic growth season climate distance (Figure S7d).

**Figure 5.**
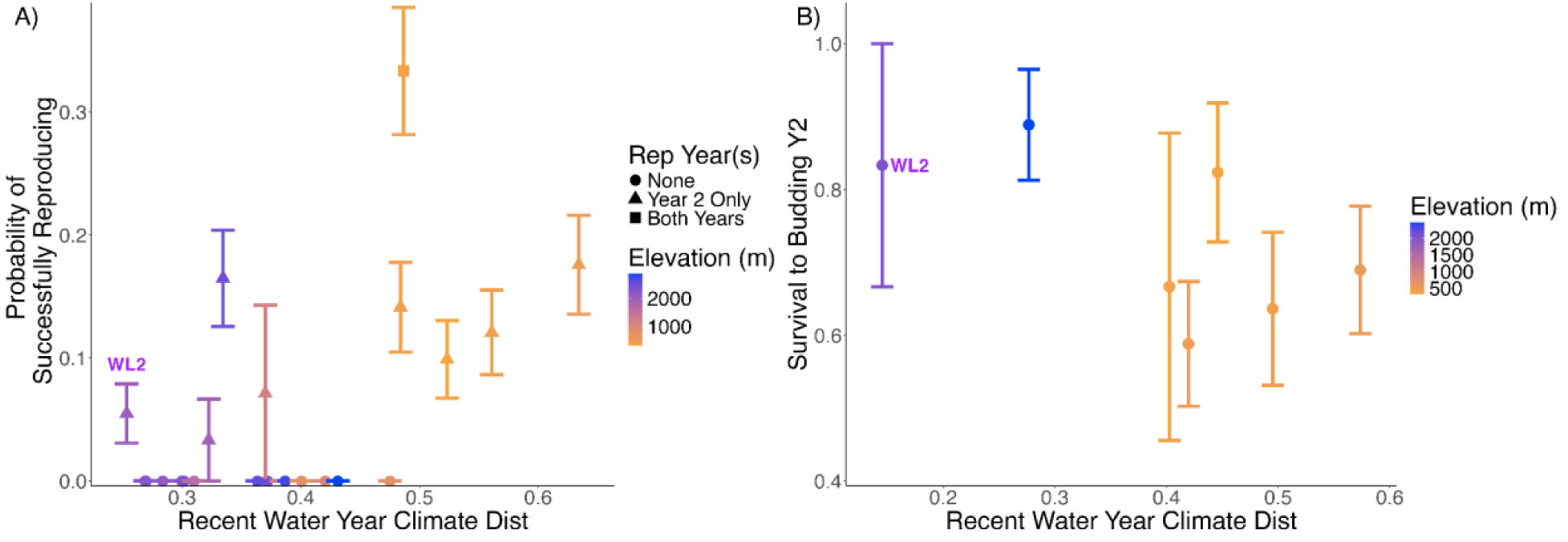
The relationship between recent water Gower’s climate distance and A) the probability of successfully reproducing in either year 1 or 2, and B) survival to budding in year 2. In A) the years in which a population had reproduction are represented by shapes. In both panels, points represent population means +/- standard error. Colors represent population elevation in meters. The native WL2 population is labeled.

**Table 3.** Results from linear and logistic regressions testing for an effect of climate and geographic distance on fitness. Population and block were included in all models as random effects. Maternal family nested within population was also included for first year and winter survival, and fruit number in year 2. Only 2 populations had greater than 1 individual survive to reproduce in year 1 so we did not analyze survival to budding and fruit number for year 1. For each trait, the effects of water year vs. growth season and recent vs. historical climate were analyzed separately, resulting in 4 models. Only the effect of the water year climate distance was tested for winter survival since the growth season does not include the winter. The two composite reproductive fitness estimates were analyzed with the average climate distance from the garden across 2023 and 2024. Coefficient estimates are shown with asterisks representing significance. P-values are represented as ***P < 0.001, **P < 0.01, *P < 0.05, ^P <0.08.

|  | Model 1 - Water Year, Recent |  | Model 2 - Water Year, Historical |  | Model 3 - Growth Season, Recent |  | Model 4 - Growth Season, Historical |  |
| --- | --- | --- | --- | --- | --- | --- | --- | --- |
|  | Climate Dist | Geo Dist | Climate Dist | Geo Dist | Climate Dist | Geo Dist | Climate Dist | Geo Dist |
| Estab. Surv | 0.31*** | -0.32*** | 0.18^ | -0.30** | -0.19^ | -0.27** | -0.29** | -0.23** |
| First Year Survival | 1.07*** | -0.27 | 0.67* | -0.22 | -0.43 | -0.18 | -0.66* | -0.09 |
| Winter Survival | 1.44** | -0.26 | 1.16* | -0.31 | N/A | N/A | N/A | N/A |
| Surv to Buds Y2 | -0.61* | 0.50* | -0.53* | 0.47^ | -0.51* | 0.25 | -0.33 | 0.29 |
| Fruit N Year 2 | -0.08 | 0.25^ | -0.06 | 0.24^ | -0.10 | 0.23^ | -0.05 | 0.23^ |
| Prob. of Fruits | 1.64* | -0.64 | 1.12^ | -0.63 | 0.10 | -0.58 | -0.84 | -0.47 |
| Total Rep Output | 0.00 | 0.04 | 0.05 | 0.02 | -0.37* | 0.16 | -0.40* | 0.14 |

Geographic distance was not a consistent predictor of fitness, after accounting for climate distance (Figure S8; Tables 2,3). Establishment was significantly lower in populations that were geographically further away from the garden (Figure S8b). Survival to budding in year 2 increased with geographic distance, although that effect was only significant in the recent water year model (Figure S8c; Table 3).

To account for the fact that some flowers had not matured into fruits at the reproductive census, we compared the analysis of fruit production to an analysis of total potential reproductive output (flowers and fruits summed). The only difference in the analyses for fruit production versus flower plus fruit production was in year 2 where the positive effect of geographic distance was significant, rather than marginal, in all flower plus fruit models (Estimates: 0.27-0.33; P-values: 0.022-0.034).

### Is there evidence for local adaptation or adaptational lag in the native population?

The above results confirm that populations of *S. tortuosus* have already started experiencing the effects of climate change. Consequently, it is important to test whether the native population is still locally adapted or if the recent climate change has resulted in maladaptation. The native WL2 population had lower establishment, first year survival, over-winter survival, and probability of successfully reproducing than some non-local populations (Figure 5a, S4, S6). In contrast, the WL2 population was not necessarily maladapted in terms of survival to budding in year 2 and total fruit number (Figure 5b; S5). There was no clear evidence of a home-site advantage in any fitness component. It is important to note that although WL2 had lower establishment and first year survival than some non-local populations, the average survival probability for both fitness components was higher than 55%.

### Do weather events within growing seasons have similar effects on fitness as climate?

To evaluate whether weekly weather had similar effects on fitness as long-term climate, we also tested the relationship between weekly mortality and soil temperature and moisture. The weekly climate results support the water year climate distance results with warmer temperatures in the preceding time period leading to lower weekly survival (Table 4). Soil temperature one week and two weeks prior to a field survey were the most predictive time periods. Increased minimum soil temperature negatively affected survival at each time point prior to the field surveys. Diurnal range and soil moisture content (VWC) did not significantly affect mortality in the field. There was a positive effect of weeks since transplant in each model as the probability of mortality increased over time.

**Table 4.** Results from logistic regressions testing for the effect of weekly climate on weekly mortality at the common garden. Each model had a fixed effect for soil temperature or moisture (VWC) at different timepoints before each field survey. The number of weeks since transplant was included as a fixed effect with population and block as random effects. Coefficient estimates are shown with asterisks representing significance. P-values are represented as ***P < 0.001, **P < 0.01, *P < 0.05, ^P <0.08.

|  | 1 Week<br>Prior to Census |  | 2 Weeks<br>Prior to Census |  | Days 6-13<br>Prior to census |  |
| --- | --- | --- | --- | --- | --- | --- |
|  | Envt | Weeks | Envt | Weeks | Envt | Weeks |
| Min Soil Temp | 0.20** | 0.33*** | 0.45*** | 0.50*** | 0.24* | 0.28*** |
| Mean Soil Temp | 0.17** | 0.32*** | 0.23** | 0.39*** | 0.04 | 0.18* |
| Max Soil Temp | 0.05* | 0.15*** | 0.03 | 0.17*** | 0.01 | 0.15** |
| Diurnal Range<br>Soil Temp | 0.04^ | 0.11* | 0.02 | 0.14*** | 0.00 | 0.14** |
| Mean Soil VWC | -5.92 | 0.18*** | -5.49 | 0.19*** | -4.36 | 0.20*** |

## Discussion

As the climate changes, understanding population and species persistence requires investigation of whether plants can adapt fast enough to track shifting local conditions and the potential success of upslope migration (Anderson & Wadgymar, 2020). This is especially important in mountains where climate gradients are steep and seasonal timing may vary significantly (Menges & Waller, 1983; Satyanti et al., 2021). Our findings provide evidence for adaptational lag at high elevation populations in a system where climate has shifted substantially over the last three decades. Low elevation populations outperform high elevation populations for several fitness components. However, we also found evidence for local adaptation at later life history stages in high elevation populations, highlighting the importance of measuring fitness across the life cycle (Satyanti et al., 2021). Therefore, higher elevation plants may be slow to adapt to the warmer conditions they may be experiencing during their first year of life, but they still have adaptations to survive and initiate reproduction in their second year. This suggests that strategic assisted gene flow that combines low elevation alleles that allow for high first year survival with high elevation alleles that facilitate reproductive success in the second year may be key for conservation management (Aitken & Whitlock, 2013; Browne et al., 2019). Some climate distance effects on fitness were only present when the entire water year was considered, as opposed to just the growing season, underscoring the need to consider factors shaping life histories and seasonal timing. Our results also highlight the importance of looking at both multidimensional and directional climate distances when evaluating fitness differences in common garden studies. Specifically, we found that some fitness components were related to temperature distance whereas others, namely winter survival, were only related to multivariate climate distance.

### Evidence for Adaptational Lag

We predicted that low elevation populations from historically warm climates would be better adapted than high elevation populations to the contemporary warmer climate in the high elevation garden site, resulting in higher fitness. In fact, high elevation populations from recent climates more similar to the 2023-2024 high elevation garden climate, including the native population at that site, had lower fitness in year 1, over-winter, and compositely than populations further away from the garden climate. This result is consistent with other studies that have found adaptational lag, or local maladaptation potentially due to climate change (Anderson & Wadgymar, 2020; DeMarche et al., 2016; Wilczek et al., 2014). *S. tortuosus* populations from high elevation may be at risk of population decline, and potentially extinction, due to this adaptation lag. As expected, if there is a directional adaptation lag, most populations from higher elevations than WL2 performed similarly or worse than the native WL2 population. Interestingly, neither the historic nor recent low elevation climate was more similar to the garden’s 2023-2024 climate than the high elevation populations’ home climates.

Furthermore, populations from sites with warmer water year climates than the 2023-2024 high elevation garden had the highest survival in year 1 and compositely. These results are consistent with the results that weeks with higher temperatures led to higher mortality rates in the garden itself. Thus, high temperature is likely a strong selective pressure for this species. Our results also show that the climate at all home sites included in this study have shifted to be warmer in the past 30 years. Other studies have found similar results with plants from warmer home sites performing better than the local population (Anderson & Wadgymar, 2020; Browne et al., 2019; Martínez-Berdeja et al., 2019; Villoutreix et al., 2025; Wilczek et al., 2014). Due to the life history differences across the range of *S. tortuosus*, populations that are warmer during the water year are also wetter during the growing season. Thus, the populations from warmer water year climates than the garden with lower survival to budding in year 2, were the same populations that came from historically wetter growth seasons and had low total fruit production. Similarly, while lower elevation populations having higher winter survival than high elevation populations may seem surprising, low elevation populations likely experience winter frost damage more frequently than high elevation populations and therefore potentially selected to be more frost tolerant. If snow comes late or melts early with climate change at high elevations, the risk of frost damage increases (Pardee et al., 2018). At high elevations where the first precipitation comes as snow plants historically may have been insulated by snow through the winter. Even though winter survival was only related to multidimensional climate distance, we did not have access to potentially more relevant variables like the frequency of frost events.

Altogether these results suggest that 1) high elevation populations have lagged in responding adaptively to increasing temperatures and 2) low elevation populations from both recent and historically warmer home sites were able to take advantage of growing in a cooler high elevation garden, particularly in the first year. An alternative hypothesis would be that some of the high elevation populations came from poor quality sites or populations with high levels of inbreeding or genetic load. Poor quality sites or high genetic load could result in a reduction of fitness independent of climate (Byers & Waller, 1999).

### Variable Climate Distance Effects on Fitness across Life Cycle Stages

Despite the positive relationship between fitness and climate distance for year 1 and over-winter survival, populations closer to the garden’s climate had higher survival to budding in year 2 and total reproductive output. Since this plant often exhibits a biennial or perennial life cycle, particularly at higher elevations, surviving to the second year of life is required to even get a chance to reproduce. Our results suggest a tradeoff between year 1 and 2 fitness components at high elevations. However, the lower survival to budding in year 2 for low elevation populations is not due to an inability to switch from an annual to biennial life history. Low to mid-elevation populations survived the winter and thus at least attempted a biennial life cycle. Interestingly, the composite fitness metric of probability of reproducing followed the trends of year 1 and over-winter survival rather than year 2 survival to budding. These results highlight the importance of measuring multiple fitness components: measuring only one, such as survival to budding in this case, or reproductive fitness, could obscure the dynamic pattern of selection across the lifespan (DeMarche et al., 2016; Mojica & Kelly, 2010).

Finding maladaptation in some fitness components and local adaptation in others could be the result of life history tradeoffs, or life history evolution in response to seasonal conditions. For example, a study on *Boechera stricta* found local adaptation in viability, but maladaptation in fecundity at a high elevation common garden site (Anderson et al., 2014). Theory predicts that this dynamic can result due to tradeoffs between fitness components (Cotto et al., 2019), however we did not find any significant tradeoffs between survival and reproduction within a year. The variation in fitness we found is more tied to early versus late portions of the life cycle, suggesting that life history and seasonal timing and conditions may be driving some of the differences we found. Assisted gene flow may be a useful conservation strategy in cases such as these. For example, assisted gene flow from low elevation to high elevation in *S. tortuosus* could introduce the traits/alleles that lead to higher survival through the first year of life while maintaining the traits/alleles already present in high elevation populations that lead to higher reproductive success in the second year of life (Aitken & Whitlock, 2013). Future studies are necessary to test whether high elevation populations have sufficient genetic variation in the traits that lead to higher survival through the first year to respond to the climatic changes they are experiencing or whether assisted gene flow should be explored.

It is important to note that these results represent one generation and using wild collected seeds. Since we used wild collected seed, the phenotypes that contributed to fitness included maternal effects, and not just within season climate effects. In addition, although we have observed low survival to the second year in many of the natural high elevation population sites (Gremer unpublished data), it is unknown whether the low elevation populations that survived better than the high elevation populations would be able to do so consistently across years. Temporal variation in the strength and direction of selection across years can be common (Siepielski et al., 2009), as we see here. In our system, year to year climate variation at the home sites is much larger than the change in mean climate from historical to recent time (Figure S2). Thus, the selective optimum probably varies from year to year potentially resulting in different populations or genotypes having higher fitness depending on the year of the study. Population demography can also vary within sites across years (Jeong et al., 2022; Morrison et al., 2022). Studies that have repeated their common garden experiment across years have found mixed results with regard to the consistency of fitness differences between populations across years (Anderson & Wadgymar, 2020; DeMarche et al., 2016). Future studies should thus replicate common garden plantings across years to evaluate the consistency of the fitness differences and evidence for adaptational lag.

Another caveat is that we forced all plants to germinate and transplanted them to the field at the same time. Thus, our study does not take into account the potential differences in germination and recruitment between populations. It is likely that if we had not experimentally forced germination timing, the low elevation populations would have germinated in the fall (Gremer, Chiono, et al., 2020; Gremer, Wilcox, et al., 2020). Young seedlings would likely not survive the winter well, potentially reducing the overall lifetime fitness of those populations. In other words, low elevation populations may be maladapted at earlier life stages than we captured in our study. In contrast, by transplanting seedlings in early summer, the faster growth rate of low elevation populations could have resulted in the higher over-winter survival we found as those plants were bigger pre-winter than high elevation plants (Quarles-Chidyagwai, *unpublished)*. More research is needed to test these hypotheses and any assisted gene flow strategies should take into account the early life stages as well.

### Water Year versus Growth Season Climate

Life history differences across elevations can act as a form of niche construction, thereby constraining the growth season climate (Clark et al., 2020; Donohue, 2005). A study across the *Streptanthus/Caulanthus* clade found that the “lived” climate niche was more constrained than the annual climate niche (Bontrager et al., 2025). Our study provides some support for this as the climate distance from the 2023 garden was higher with the climate summarized by water year than by growth year. In addition, the relationship between Gower’s climate distance and establishment and first year survival reversed between the water year and growth season. This result is likely due to the fact that the low elevation populations’ historic growth seasons were more similar to the garden than their recent and historic water years. Despite the likelihood of niche construction in this species, the water year climate is still important because it helps determine the right time of year to grow (Giesel, 1976; Varpe, 2017). Thus, the water year can influence both fitness components that determine seasonal transitions, like over-winter survival, and fitness components that occur during the growth season. Studies that investigate species with a wide range of life history strategies should carefully consider what climate summary to use when investigating adaptive differentiation and fitness trends across climatic gradients (Evers et al., 2021).

## Conclusions

Our study contributes to the growing evidence that plants that grow in high elevation environments may be at risk of population decline due to climate change. Populations from lower elevations are expected to be pre-adapted to the warmer conditions that high elevations are projected to experience with climate change and thus may provide an avenue for evolutionary rescue. However, seasonal conditions and timing, and consequently plant life history can vary across elevations, such that low elevation populations are only partially adapted to contemporary and future high elevations. These differences in seasonality and life history can have important consequences on climate adaptation and fitness components across the life cycle can vary in the direction of the response to climate. It is important to consider this complexity when evaluating population persistence in the face of climate change and possible scenarios of evolutionary rescue from upslope dispersal or gene flow.

## Acknowledgments

This work benefited greatly from discussions with Sharon Strauss, Denneal Jamison-McClung, and Troy Magney. In addition, the authors thank Michelle Stern at the USGS for assistance with the climatic variables. We also acknowledge postdoctoral scholar, Rishav Ray, graduate student Maya Arakaki, technician Kate Ruiz-Cox, and numerous undergraduates, including Mia Ashby, Sophie Benefiel, Eda Ceviker, Christina Chen, Kate DiTrani, Bryce Johnson, Hugo Mahatdejkul, Victoria Mattson, Katie Michaels, Samantha Swan, Megan Wong, and Sammie Yee, for help with the experiment and discussion of results. This research was funded by the National Science Foundation (DEB-2129589 to J.N.M and J.R.G, DEB-1831913 to J.R.G, J.N.M., J. Schmitt and S.Y. Strauss) and the National Institute of Food and Agriculture award (CA-D-PLB-2795-H to J.N.M). This work took place within the ancestral territories of the Miwok, Washoe and Nisenan people.

## Author Contributions

J.R.G, J.N.M, and J.S. conceptualized and designed the study. B.Q-C and S.R.A performed the experiment and gathered the data with assistance from all authors. B.Q-C, J.R.G, and J.N.M designed the data analysis. B.Q-C and J.N.M analyzed the data with assistance from J.R.G. J.R.G. and J.N.M contributed equally to this manuscript. B.Q-C wrote the manuscript with contributions from all other authors. All authors contributed critically to the development of ideas, analyses, and interpretation of results. **Statement on inclusion:** The study species is native to California, the work took place in California, and all authors live in this state. We presented results from this work to stakeholders within the region of the study (J.R.G and J.N.M to Tuleyome and B.Q-C to California Native Plant Society). We also incorporated undergraduate students from California into our field survey team and taught them data analysis skills with some of the data. The exact site of the common garden was chosen to avoid places of importance to First Nations people.

## Data Availability Statement

Data will be posted on Zenodo and GitHub upon publication.

