## Supplementary Figures and Tables for "Adaptational lag at high elevations depends on life stage in a California wildflower"

**Table S1.** Home sites included in the common garden experiment with year of seed collection, location in latitude, longitude, and elevation. The number of maternal families for each population at transplant is listed, along with the number of plants per family. Sites that received less than 70mm of average annual snowpack are denoted with a "^" and populations that received greater than 70mm of average annual snowpack are denoted with a "*".

| Site name | Code | Seed year | Latitude | Longitude | Elevation (m a.s.l) | Maternal Family N | N plants per family |
|---|---|---|---|---|---|---|---|
| Canyon Creek^ | CC | 2018 | 39.58597 | -121.43311 | 313 | 7 | 13 |
| Table Mountain 2^ | TM2 | 2021 | 39.59255 | -121.55072 | 379 | 7 | 4-22 |
| Sutter Creek^ | SC | 2021 | 38.41167 | -120.73583 | 421 | 6 | 1-27 |
| Iowa Hill^ | IH | 2021 | 39.09332 | -120.92114 | 454 | 7 | 13-14 |
| Ben Hur^ | BH | 2021 | 37.40985 | -119.96458 | 511 | 7 | 13 |
| Weaverville* | WV | 2014 | 40.74084 | -123.00364 | 749 | 2 | 1-2 |
| Feather River/ Rich Bar* | FR | 2022 | 40.01362 | -121.18498 | 787 | 7 | 2-13 |
| Drum Powerhouse Road* | DPR | 2020 | 39.22846 | -120.81518 | 1019 | 7 | 3-21 |
| Washington Road* | WR | 2016 | 39.34346 | -120.80659 | 1158 | 3 | 1-11 |
| Wright's Lake 1* | WL1 | 2020 | 38.78608 | -120.2143 | 1614 | 6 | 3-21 |
| Sequoia 1* | SQ1 | 2021 | 36.56435 | -118.7764 | 1921 | 6 | 2-10 |
| Sequoia 2* | SQ2 | 2021 | 36.66557 | -118.83543 | 1934 | 7 | 1-27 |
| Wright's Lake 2* | WL2 | 2020 | 38.8263 | -120.25242 | 2020 | 7 | 13 |
| Yosemite 4* | YO4 | 2021 | 37.76378 | -119.77244 | 2158 | 7 | 1-15 |
| Carson Pass 2* | CP2 | 2022 | 38.66169 | -120.13065 | 2244 | 7 | 6-15 |
| Carson Pass 3* | CP3 | 2022 | 38.70649 | -120.08797 | 2266 | 7 | 7-15 |
| Lassen Volcanic 3* | LV3 | 2016 | 40.47917 | -121.52311 | 2354 | 6 | 1-13 |
| Sequoia 3* | SQ3 | 2021 | 36.72109 | -118.84933 | 2373 | 7 | 2-7 |
| Yosemite 7* | YO7 | 2022 | 37.80903 | -119.56605 | 2470 | 7 | 4-26 |
| Yosemite 8* | YO8 | 2021 | 37.81117 | -119.48624 | 2591 | 7 | 9-14 |
| Lassen Volcanic 1* | LV1 | 2018 | 40.47471 | -121.50486 | 2593 | 7 | 12-14 |
| Lassen Volcanic Trail 1* | LVTR1 | 2020 | 40.47917 | -121.5032 | 2741 | 7 | 6-23 |
| Yosemite 11* | YO11 | 2021 | 37.93844 | -119.23157 | 2872 | 7 | 4-18 |

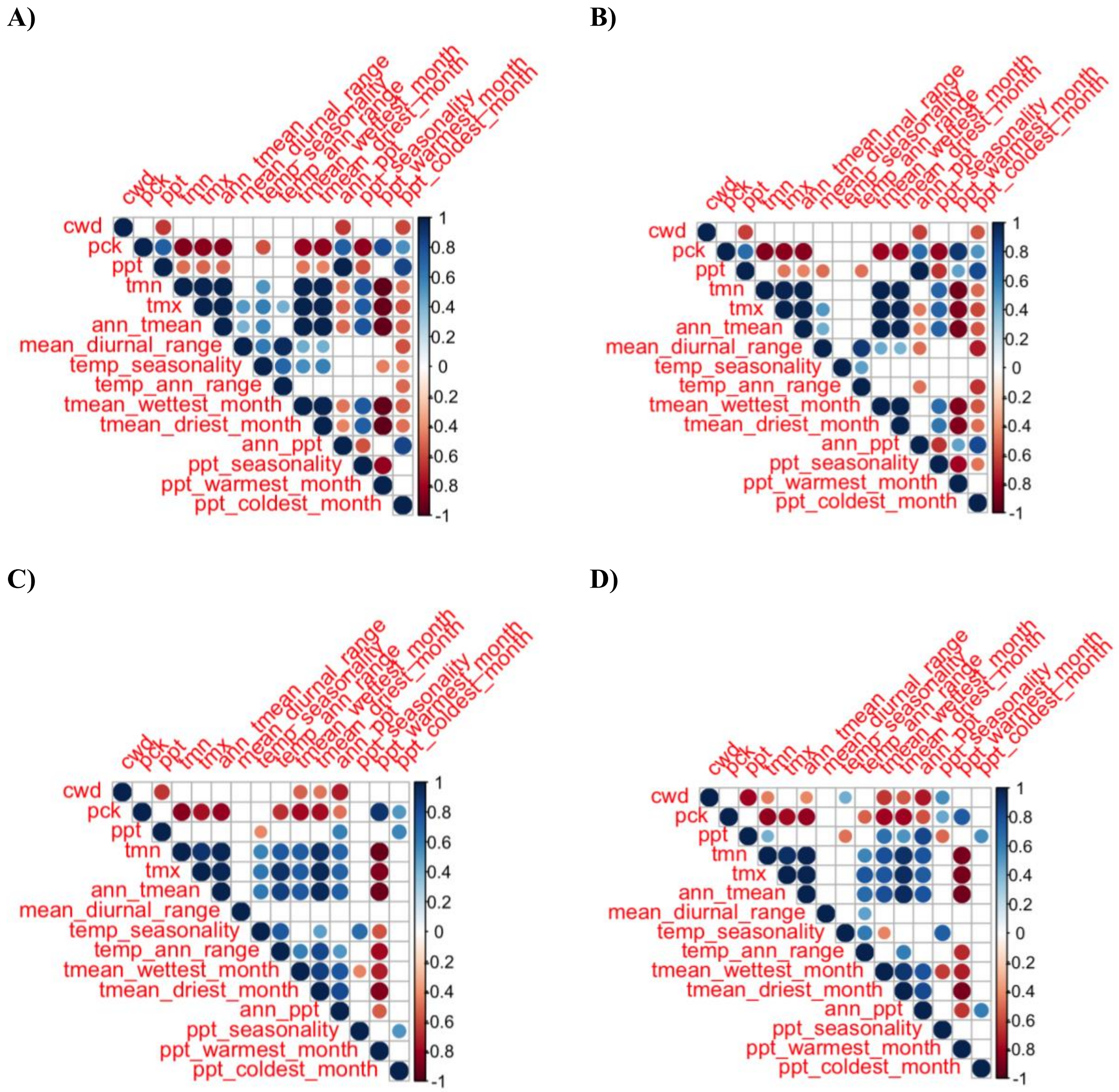


**Figure S1.** Pearson's correlations between climate variables that were used to compute Gower's climate distance. (A-B) water year, historic and recent, (C-D) growth season, historic and recent. The historic time period (1964-1993) is on the left and the recent time period (1994-2023) is on the right. Non-significant correlations are blank.

**Table S2.** Loadings from principal components analysis of the average climate at all sites. Water year, growth season, and recent historical time periods are included. TMN and TMX were excluded from the PCA due to being highly correlated with ann_tmean (see Fig. S1).

| Variable | PC1 | PC2 | PC3 | PC4 |
|---|---|---|---|---|
| CWD | 0.14 | -0.33 | -0.37 | -0.10 |
| PCK | -0.40 | 0.12 | -0.15 | -0.18 |
| PPT | -0.30 | 0.32 | 0.24 | 0.05 |
| Ann_Tmean | 0.42 | 0.17 | 0.11 | 0.10 |
| Mean Diurnal Range | 0.12 | -0.09 | 0.01 | -0.63 |
| Temp_Seasonality | 0.07 | 0.26 | -0.59 | -0.03 |
| Temp_Ann_Range | 0.08 | 0.36 | -0.33 | -0.44 |
| Tmean Wettest Month | 0.38 | 0.17 | 0.32 | -0.01 |
| Tmean Driest Month | 0.35 | 0.33 | 0.12 | 0.01 |
| Ann_PPT | -0.24 | 0.45 | -0.01 | -0.06 |
| PPT_Seasonality | 0.20 | -0.11 | -0.43 | 0.42 |
| PPT Warmest Month | -0.31 | -0.34 | 0.09 | -0.11 |
| PPT Coldest Month | -0.26 | 0.25 | -0.12 | 0.40 |
| Proportion of Variance | 0.36 | 0.26 | 0.15 | 0.11 |

**Table S3.** Results from A) permanovas and B) linear models testing for differences between recent and historical climate in PC space. The PCA shown in Fig. 3 and Table S2 was tested. All PCs were tested but only PCs 1-4 are shown. F values are reported with asterisks representing significance. P-values are represented as ***P < 0.001, **P < 0.01, *P < 0.05, ^P <0.08.

| | **A) Permanova Results** | **B) Linear Models of Select PC Axes** | | | |
|---|---|---|---|---|---|
| | | PC1 | PC2 | PC3 | PC4 |
| Season | 38.98** | 105.39*** | 125.10*** | 12.87*** | 7.99** |
| TimePd | 10.56** | 12.72*** | 6.33* | 10.71** | 19.98*** |
| Elevation | 76.03** | 498.37*** | 115.26*** | 18.31*** | 2.67 |
| Latitude | 22.63** | 96.23*** | 22.22*** | 7.10** | 7.04** |
| Season*TimePd | 0.74 | 0.28 | 0.18 | 0.67 | 0.51 |
| Season*Elev | 22.23** | 133.65*** | 37.42*** | 2.28 | 0.34 |
| TimePd*Elev | 2.04 | 5.78* | 0.61 | 4.50* | 0.03 |
| Season*Lat | 3.39* | 4.84* | 5.87* | 0.96 | 0.02 |
| TimePd*Lat | 0.72 | 0.05 | 0.02 | 0.05 | 1.81 |
| Elev*Lat | 4.71** | 5.9* | 0.00 | 11.43** | 0.37 |
| Season*TimePd*Elev | 0.92 | 1.10 | 0.04 | 1.71 | 0.09 |
| Season*TimePd*Lat | 0.47 | 0.88 | 0.19 | 0.51 | 0.01 |
| Season*Elev*Lat | 2.59* | 1.20 | 1.19 | 5.58* | 0.08 |
| TimePd*Elev*Lat | 0.13 | 0.43 | 0.03 | 0.04 | 0.12 |
| 4-way interaction | 0.29 | 0.87 | 0.16 | 0.26 | 0.00 |
| Overall Model | 12.43** | | | | |

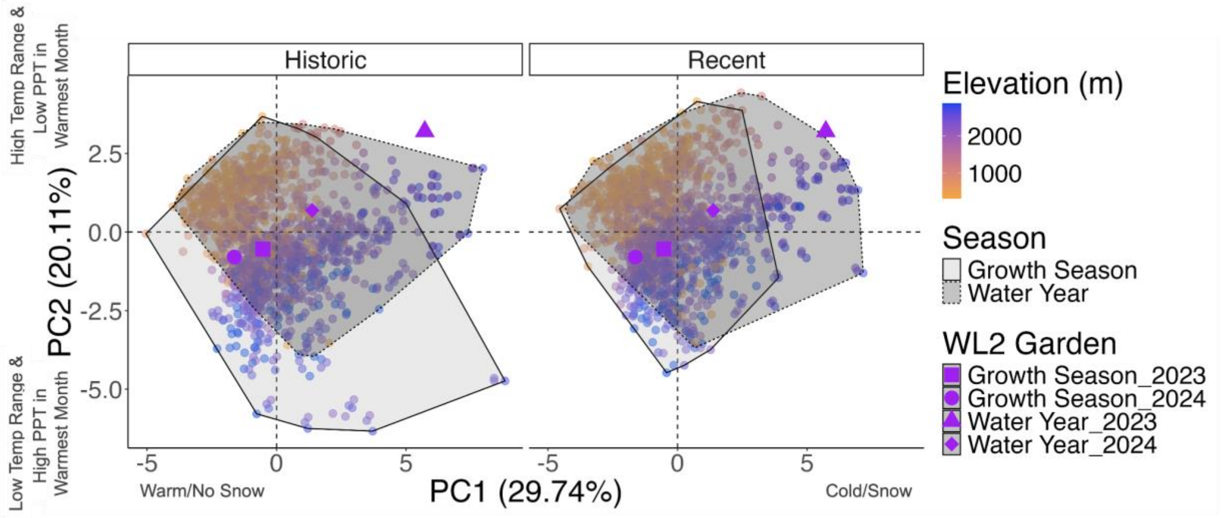


**Figure S2.** PCAs representing yearly variation in climate at all sites. The same PCA in Fig. 1A is parsed for both timeframe and seasonal summary. The historic timeframe (1964-1993) is on the left and the recent timeframe (1994-2023) is on the right. The growth season and water year are outlined by solid or dotted hulls, respectively. The garden climate during the years of this study is represented by different bright purple shapes.

**A) 2023 Water Year**

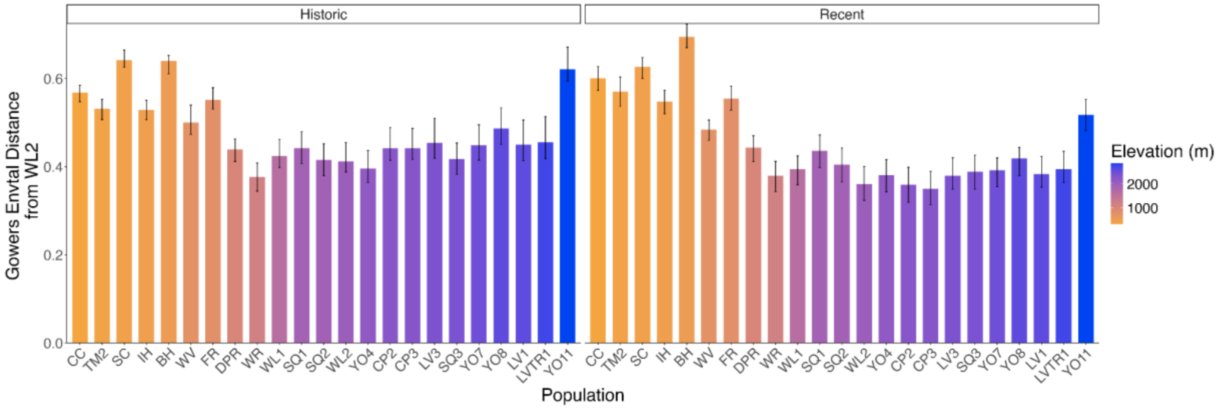


**B) 2023 Growth Season**

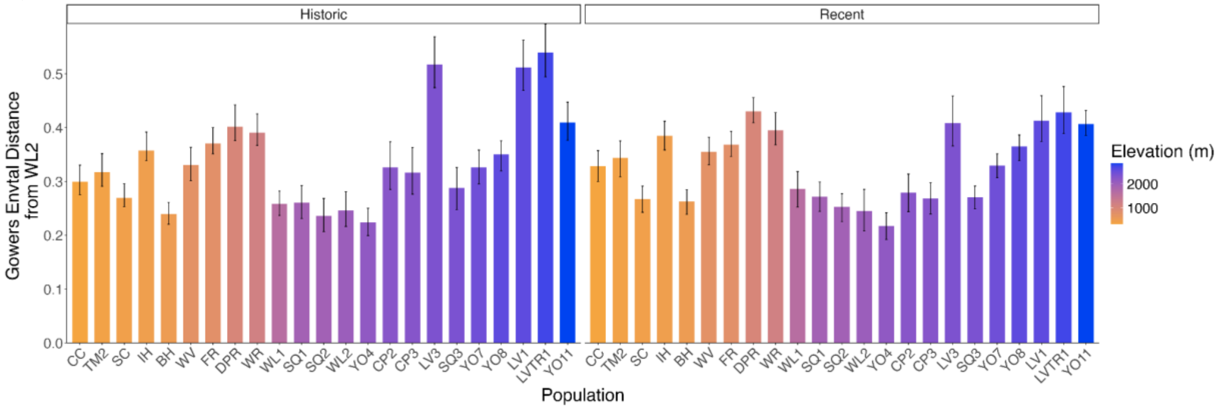


**C) 2024 Water Year**

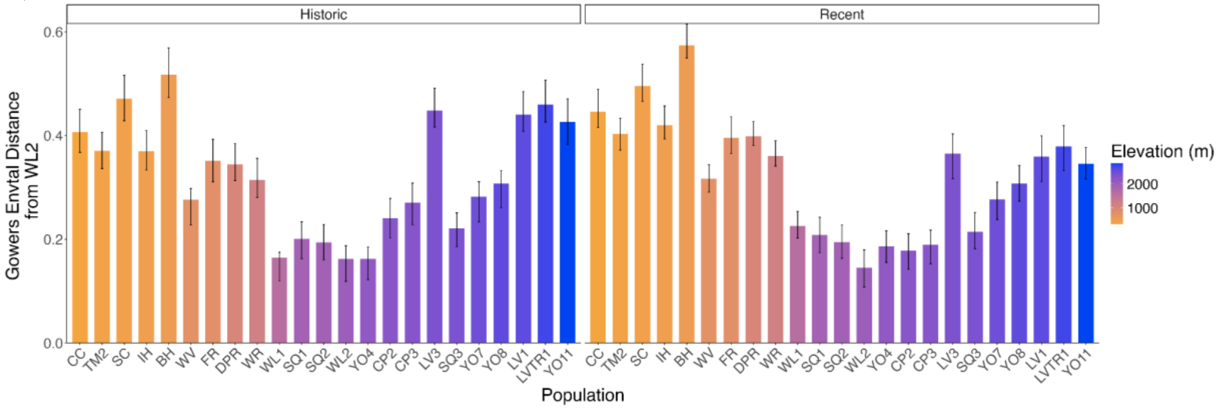

**D) 2024 Growth Season**

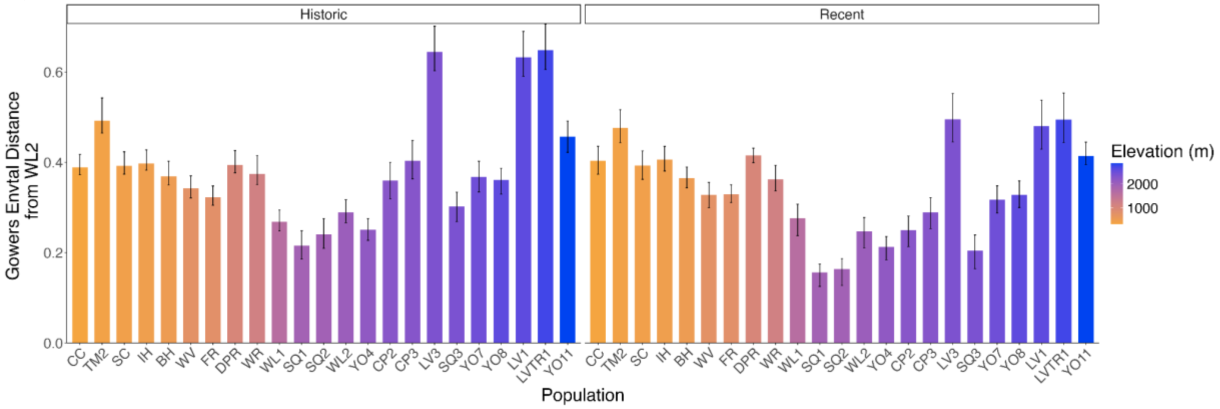


**E) Average Water Year**

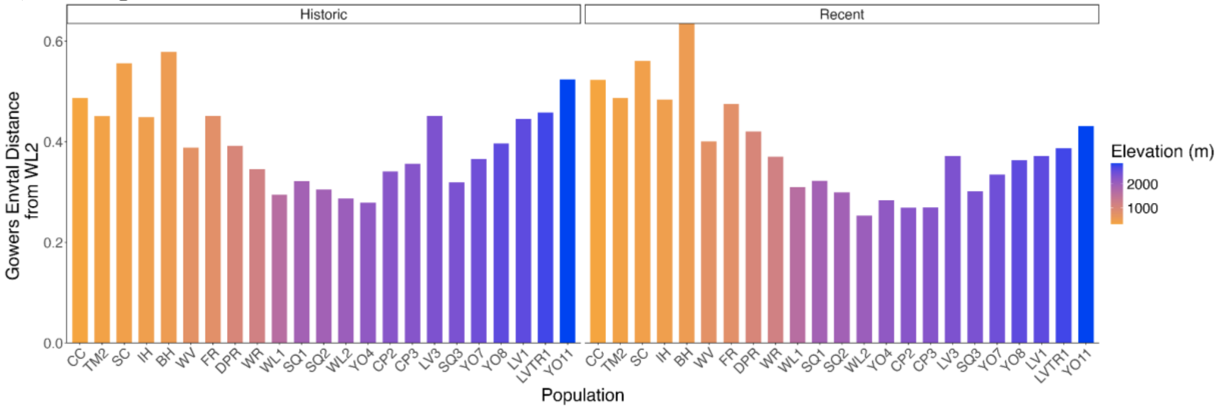


**F) Average Growth Season**

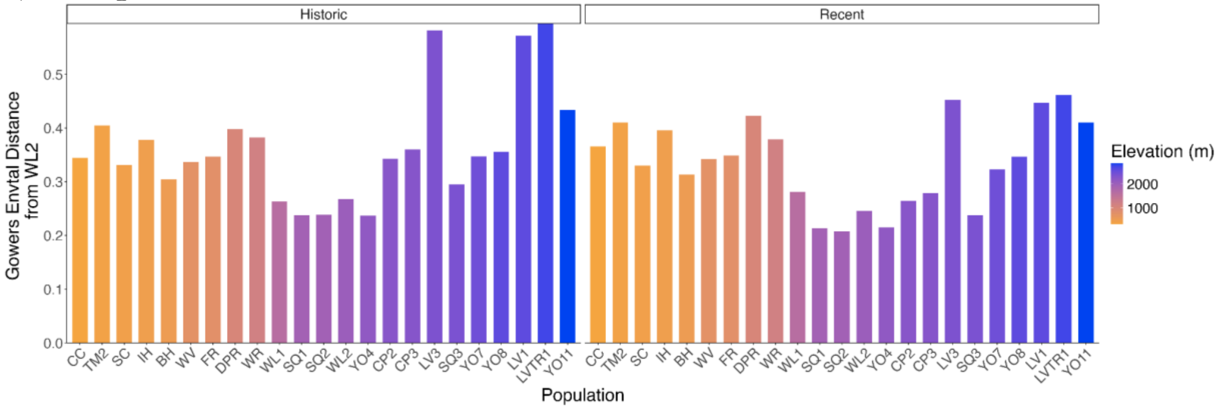


**Figure S3.** Gower's climate distance between the home sites and the garden site. Home sites' water year (A) and growth season (B) climate distance from the garden in 2023. Home sites' water year (C) and growth season (D) climate distance from the garden in 2024. The average climate distance between the home sites and the garden for the water year (E) and growth season (F). Populations are ordered on the x-axis by elevation (low to high). Colors represent home site elevation. Error bars are bootstrapped 95% confidence intervals.

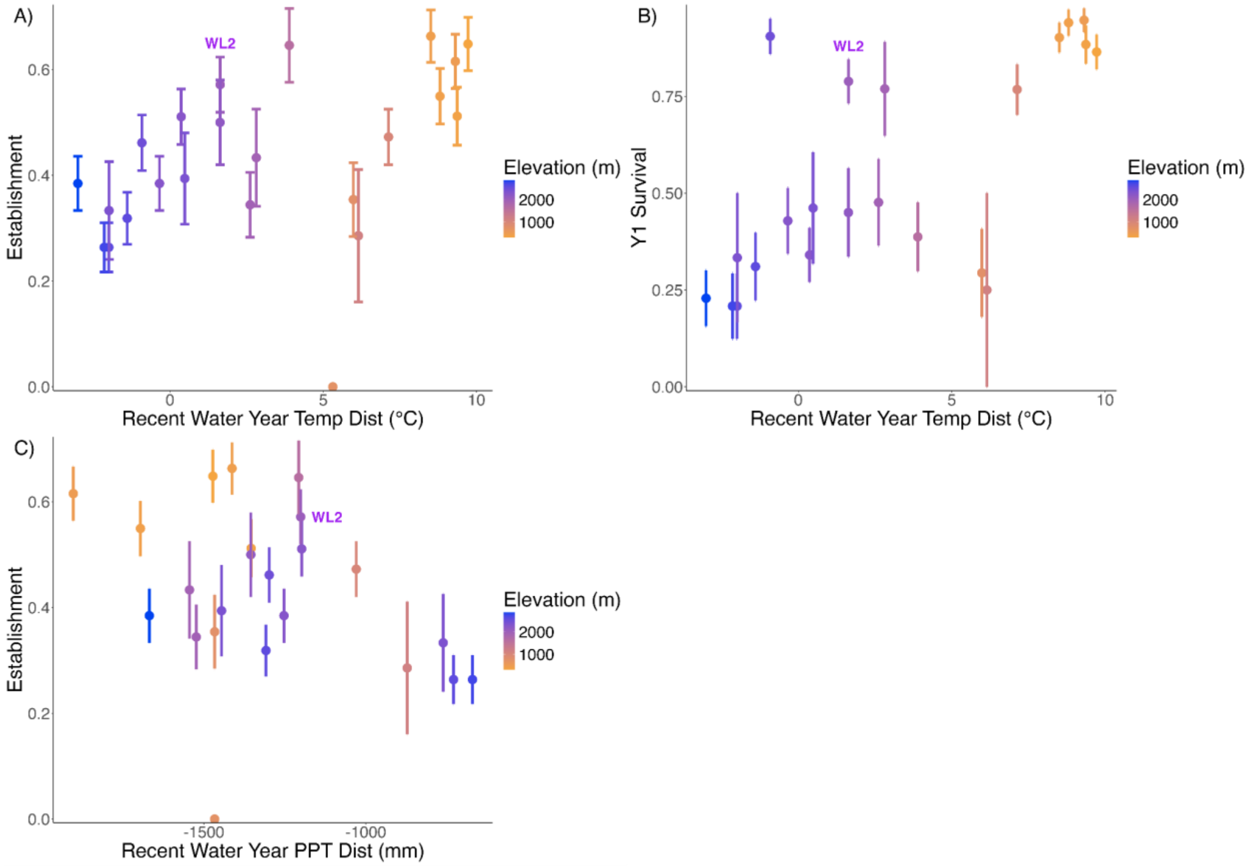


**Figure S4.** The effects of (A-B) recent water year directional annual mean temperature (BIO1) distance from home, and C) recent water year annual precipitation sum (BIO12) distance from home on establishment and first year survival. Positive values for temperature distance represent warmer temperatures at home than at the garden site and negative values for precipitation distance represent drier home sites than the garden. Points represent population means +/- standard error. Colors represent population elevation in meters. The native WL2 population is labeled.

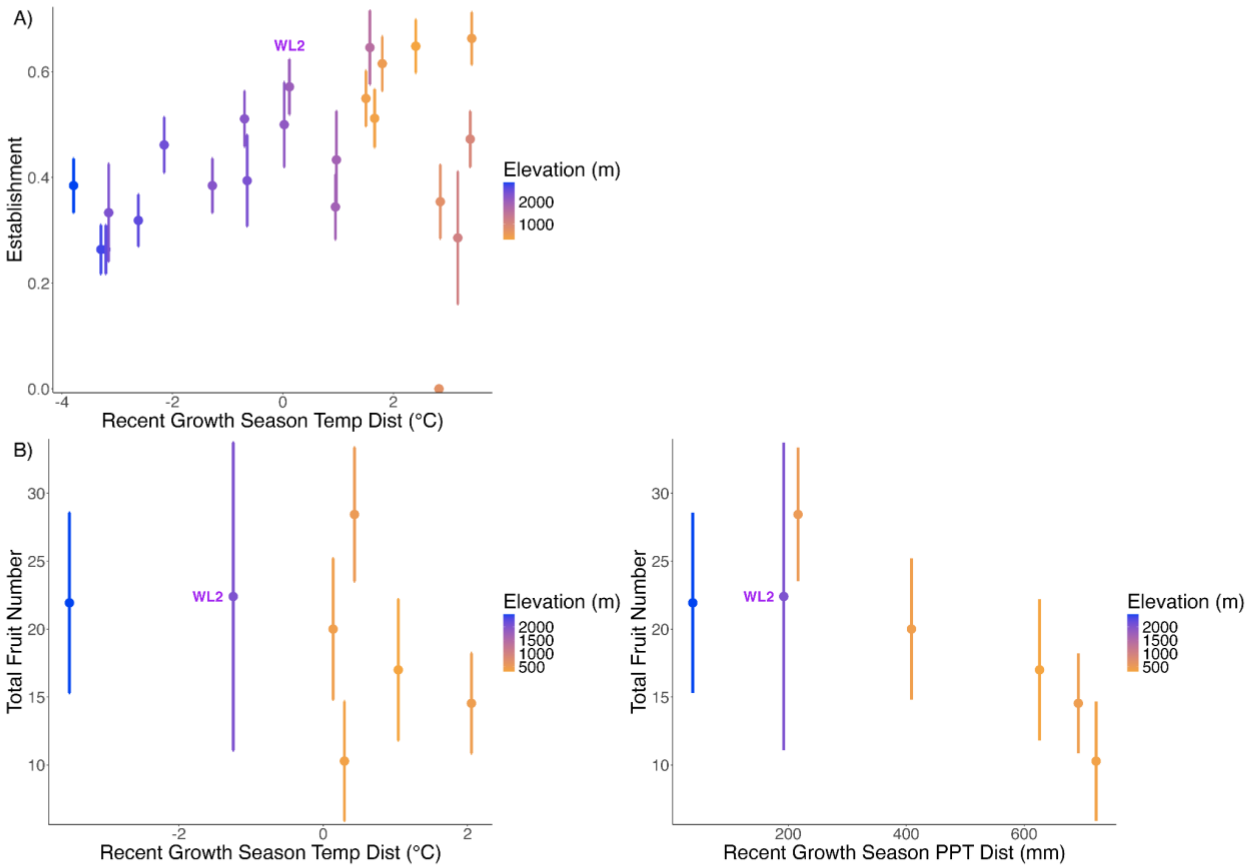


**Figure S5.** Significant growth season temperature and precipitation effects. A) The effect of recent growth season temperature distance on establishment. B) The effect of recent growth season temperature (left) and precipitation (right) distance on total reproductive output. Positive values for temperature distance represent warmer temperatures at home than at the garden site and negative values for precipitation distance represent drier home sites than the garden. In all figures, points represent population means +/- standard error. Colors represent population elevation in meters. The native WL2 population is labeled.

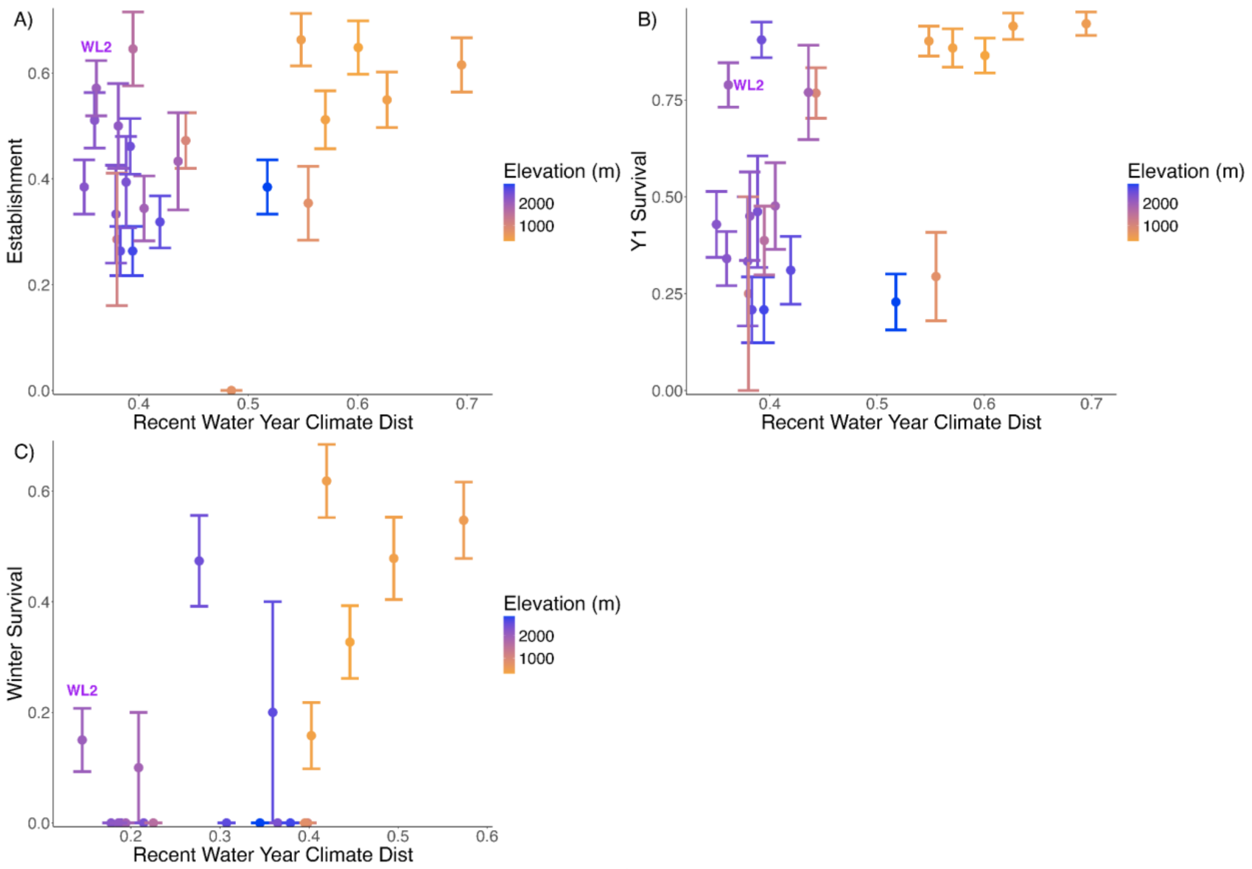


**Figure S6.** The effects of recent water year Gower's climate distance on A) establishment, B) first year survival, and C) over-winter survival. Points represent population means +/- standard error. Colors represent population elevation in meters. The native WL2 population is labeled.

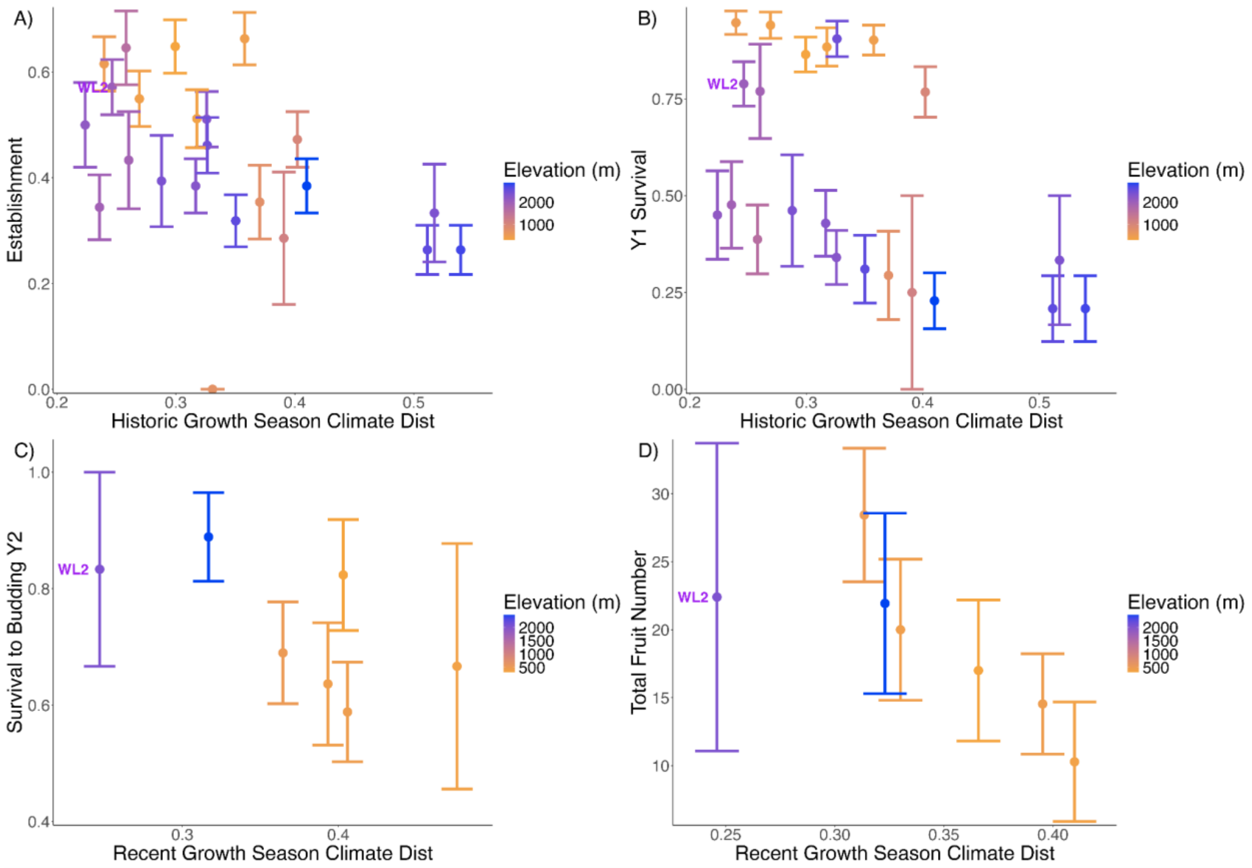


**Figure S7.** Significant growth season Gower's climate distance effects. The effect of historic growth season climate distance on A) establishment and B) first year survival. The effect of recent growth season climate distance on C) survival to budding in year 2 and D) total reproductive output. Points represent population means +/- standard error. Colors represent population elevation in meters. The native WL2 population is labeled.

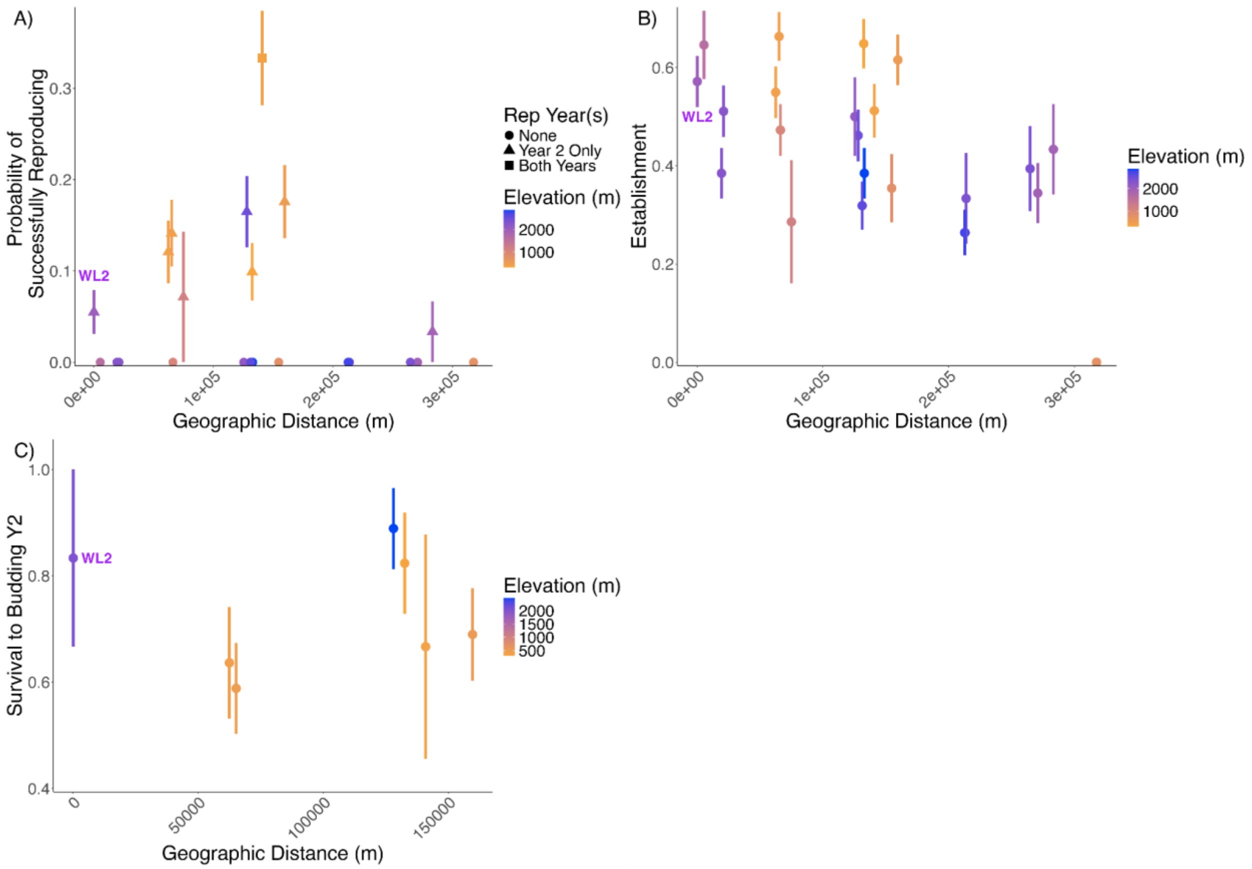


**Figure S8.** The relationship between geographic distance and A) the probability of successfully reproducing in either year 1 or 2, B) establishment, and C) survival to budding in year 2. In A) the years in which a population had reproduction are represented by shapes. In all panels, points represent population means +/- standard error. Colors represent population elevation in meters. The native WL2 population is labeled. Statistics can be found in Tables 2 and 3.
